# Increased hyaluronan by naked mole-rat HAS2 extends lifespan in mice

**DOI:** 10.1101/2023.05.04.539405

**Authors:** Zhihui Zhang, Xiao Tian, J. Yuyang Lu, Kathryn Boit, Julia Ablaeva, Frances Tolibzoda Zakusilo, Stephan Emmrich, Denis Firsanov, Elena Rydkina, Seyed Ali Biashad, Quan Lu, Alexander Tyshkovskiy, Vadim N. Gladyshev, Steve Horvath, Andrei Seluanov, Vera Gorbunova

## Abstract

**Abundant high molecular weight hyaluronic acid (HMW-HA) contributes to cancer resistance and possibly longevity of the longest-lived rodent, the naked mole-rat**^1, 2^**. To study whether the benefits of HMW-HA could be transferred to other animal species, we generated a transgenic mouse overexpressing naked mole-rat hyaluronic acid synthase 2 gene (nmrHAS2). nmrHAS2 mice showed increase in hyaluronan levels in several tissues, and lower incidence of spontaneous and induced cancer, extended lifespan and improved healthspan. The transcriptome signature of nmrHAS2 mice shifted towards that of longer-lived species. The most striking change observed in nmrHAS2 mice was attenuated inflammation across multiple tissues. HMW-HA reduced inflammation via several pathways including direct immunoregulatory effect on immune cells, protection from oxidative stress, and improved gut barrier function during aging. These findings demonstrate that the longevity mechanism that evolved in the naked mole-rat can be exported to other species, and open new avenues for using HMW-HA to improve lifespan and healthspan.**

## Main

Naked mole-rats are mouse-size rodents that display exceptional longevity, with the maximum lifespan of over 40 years^3^. Naked mole-rats are protected from multiple age-related diseases including neurodegeneration, cardiovascular diseases, arthritis, and cancer^4, 5^. These characteristics suggest that naked mole-rats have evolved efficient anti-aging and anti-cancer defenses. Previously, we have identified a novel anticancer mechanism in the naked mole-rat named early contact inhibition^1^ mediated by abundant high molecular weight hyaluronic acid (HMW-HA)^2^. Naked mole-rat tissues are highly enriched for HMW-HA compared to mouse and human tissues. Depleting HMW-HA from naked mole-rat cells increases their susceptibility to tumor formation in mouse xenograft models^2^.

Hyaluronan is a non-protein component of the extracellular matrix that affects biomechanical properties of tissues and interacts with cell receptors. While hyaluronan structure is deceptively simple, and conserved across kingdoms of life, with repeating disaccharide chains of N-acetyl-glucosamine and glucuronic acid, its biology is highly complex^6, 7^. The length of hyaluronan can range from an oligomer to an extremely long-form up to millions of Daltons, and the biological functions of hyaluronan depends on its molecular weight. Low molecular weight HA (LMW-HA) is associated with inflammation, tissue injury, and cancer metastasis^8–11^, while high molecular weight HA (HMW-HA) improves tissue homeostasis^12^, shows anti-inflammatory^13, 14^ and antioxidant properties^15^. The HMW-HA (>6.1 MDa) produced by naked mole-rats has unique cytoprotective properties^16^. The hyaluronan content is determined by the balance of hyaluronan synthesis by hyaluronan synthases and the breakdown of hyaluronan either by hyaluronidases or oxidative and nitrative stresses^17^. Three synthases, HAS1, HAS2, and HAS3, are responsible for producing HA. HAS2 mainly produces HMW-HA and shows higher expression in naked mole-rats compared to mice and humans^2^. Naked mole-rat tissues also show lower activity of hyaluronidases resulting in the massive accumulation of HMW-HA^2^. Age-related sterile inflammation has emerged as an important driving force of aging and age-related diseases^18, 19^. Hence anti-inflammatory functions of HMW-HA may confer anti-aging effects.

To test whether anti-cancer and potential anti-aging effects of HMW-HA can be recapitulated in species other than the naked mole-rat, we generated a mouse model overexpressing naked mole-rat hyaluronan synthase 2 gene (nmrHAS2 mouse). The nmrHAS2 mice showed higher resistance to spontaneous tumors and DMBA/TPA-induced skin cancer. Furthermore, the nmrHAS2 mice showed improved healthspan and extended median and maximum lifespan. Aged nmrHAS2 mice displayed reduced inflammation in multiple tissues and maintained better intestinal barrier function and healthier gut microbiome. In addition, accumulation of HMW-HA in tissues alleviated the activation of immune response both *in vitro* and *in vivo*. Our results suggest that HMW-HA improves healthspan by reducing inflammation due to its direct immunomodulating effect and improved intestinal barrier function. These findings open new avenues for using the longevity mechanisms that evolved in the naked mole-rat to benefit human health.

### Generation of nmrHAS2 mice

Since naked mole-rats accumulate HMW-HA in the majority of their tissues, to recreate this phenotype in mice, we chose to express naked mole-rat HAS2 gene (nmrHAS2) in mice using a ubiquitous CAG promoter. In the naked mole-rat, HMW-HA begins to accumulate postnatally^1^ as it is not compatible with rapid cell proliferation required during embryogenesis. Therefore, we controlled nmrHAS2 expression temporally using a Lox-STOP cassette. The cassette containing the nmrHAS2 gene was randomly integrated into mouse oocytes. Mice heterozygous for nmrHAS2 transgene were bred to homozygous R26-CreER^T2^ mice. The F1 progeny containing both nmrHAS2 and R26-CreER^T2^ transgenes was compared to littermates containing R26-CreER^T2^ (CreER) only (Fig. 1a). Expression of the nmrHAS2 gene was induced by injections of tamoxifen at 3 months of age (Fig. 1a). Both nmrHAS2 and control mice received tamoxifen injections.

**Figure 1.**
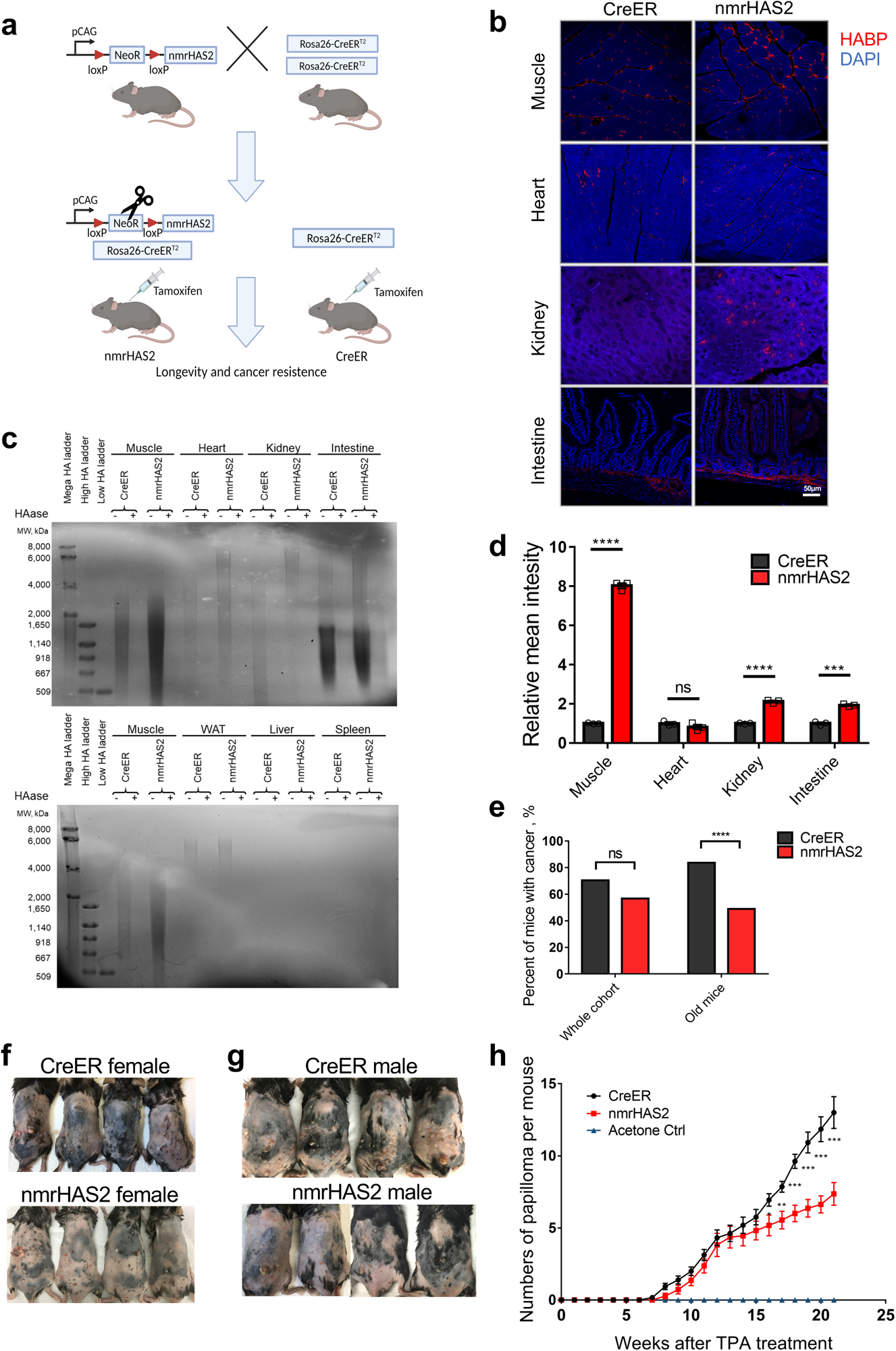
Transgenic mice overexpressing nmrHAS2 gene are resistant to both spontaneous and induced cancer. **a.** The breeding strategy of nmrHAS2 mice. Mice heterozygous with nmrHAS2 transgene were bred with mice homozygous for Rosa26CreERT2 to obtain double heterozygous nmrHAS2; CreER progeny. Single heterozygous CreER progeny was used as a control. Tamoxifen was injected when in 2-3 months animals to induce the nmrHAS2 expression. CreER mice received the same dose of tamoxifen at the same age. **b.** Representative pictures of HABP staining in multiple organs of female mice. **c.** Pulse field gel shows that female nmrHAS2 mice have higher molecular weight and more abundant hyaluronic acid in multiple tissues. HA was extracted from 200 mg of pooled tissue from two individuals. HAase treated samples were run in parallel to confirm the specificity of HA staining. The HA from muscle was loaded on both gels as a cross reference. **d.** Levels of relative HABP fluorescence intensity. N=3. *** p < 0.001, **** p < 0.0001 by unpaired Student’s t-test. **e.** Old nmrHAS2 mice (N=74) have much lower spontaneous cancer incidence compared to CreER mice (N=81). Pooled females and males. Old mice are older than 27 months. *** indicates p<0.0001, Chi-square test. **f.** Representative pictures of female mice after 20 weeks of DMBA/TPA treatment. **g.** Representative pictures of male mice after 20 weeks of DMBA/TPA treatment. **h.** nmrHAS2 mice are more resistant to DMBA/TPA induced skin papilloma. Pooled females and males. N= 7-13. ** indicates p<0.01, *** indicates p<0.001, two-tailed Student’s t-test.

Overexpression of nmrHAS2 mRNA was detected in multiple tissues of nmrHAS2 mice (Extended Data Table 1). HABP staining showed a stronger hyaluronan signal in muscle, kidney and intestines of both male and female nmrHAS2 mice compared to controls (Fig. 1b-d, Extended Data Fig. 1a-c). In addition, pulse field gel electrophoresis showed that hyaluronan extracted from the tissues of nmrHAS2 mice was more abundant and had a higher molecular weight in muscle, heart, kidney, and small intestine (Fig. 1c, Extended Data Fig. 1b). Hyaluronan levels in the liver and spleen were very low, which is consistent with these tissues being the sites of hyaluronan breakdown^20^. Interestingly, despite the high nmrHAS2 mRNA levels in most mouse organs, we observed only a mild increase in hyaluronan, likely due to high hyaluronidase activity in mouse tissues compared to the naked mole-rat^2^.

### nmrHAS2 mice are resistant to both spontaneous and induced cancer

To examine the effect of HMW-HA on lifespan and spontaneous cancer incidence, we set up aging cohorts of 80-90 mice of both genotypes. To determine the cause of death, all aged mice were checked daily, and necropsies were performed within 24 h of death. The majority of mice died from cancer, which is consistent with earlier reports that lymphomas are the common endpoint for aged C57BL/6 mice^21^. nmrHAS2 mice showed lower spontaneous cancer incidence, with 57% of nmrHAS2 mice dying from cancer compared to 70% for the control CreER group (Fig. 1e). This difference was further amplified for the oldest age group. For mice older than 27 months, cancer incidence was 83% for CreER mice and 49% for nmrHAS2 mice (Fig. 1e). This phenotype was the same across both sexes (Extended Data Fig. 1d and 1e).

nmrHAS2 mice showed accumulation of HA in the skin (Extended Data Fig. 1f.). To test the resistance of nmrHAS2 mice to chemically induced skin tumorigenesis, we treated young mice with DMBA/TPA. Sixteen weeks after TPA treatment, nmrHAS2 mice formed significantly fewer papillomas than age matched controls for females, males, and two sexes combined (Fig. 1f-h, Extended Data Fig. 1g, h). These results indicate that production of HMW-HA protected mice from both spontaneous and induced cancer.

### nmrHAS2 mice have increased lifespan

Mice of both genotypes had a similar body weight throughout their life (Fig. 2a). nmrHAS2 mice showed an increase of 4.4% in median lifespan and 12.2% in maximum lifespan (Fig. 2b). The lifespan increase was different for each sex. While females showed a more prominent 9% increase in median lifespan (Extended Data Fig. 2a), males showed a more prominent 16% increase in the maximum lifespan (Extended Data Fig. 2b).

**Figure 2.**
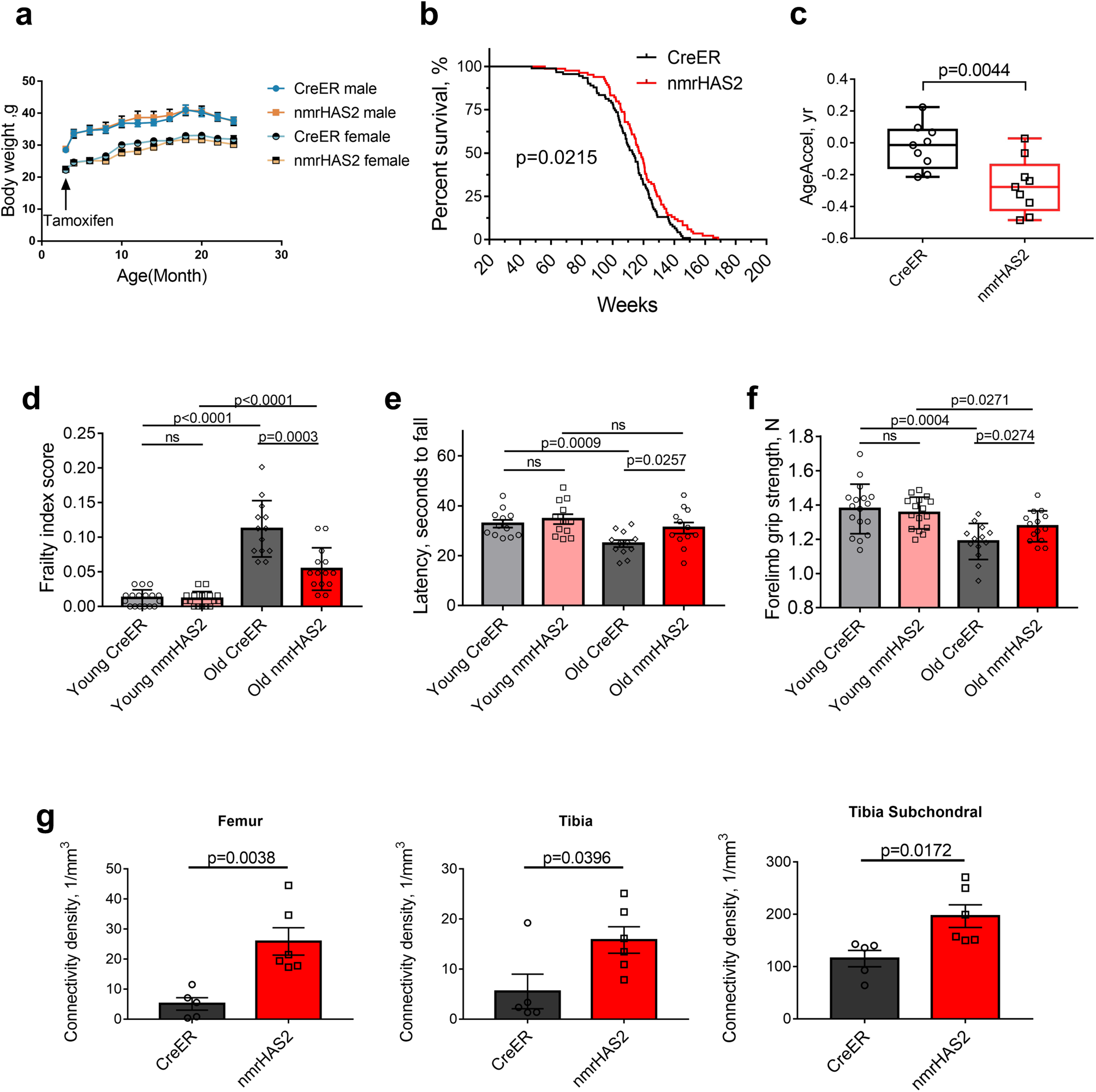
nmrHAS2 mice have extended lifespan and health span. **a.** Overexpression of nmrHAS2 gene did not affect the body weight of mice (N=9-11). The body weight of mice was measured before Tamoxifen injection, then once every month until mice reached 24 months of age. **b.** nmrHAS2 mice (N=84) had an extended median and maximum lifespan compared to CreER mice (N=91). Pooled females and males. p=0.022, log-rank test. **c.** Old nmrHAS2 mice display younger biological age. Liver DNA from 24 months old nmrHAS2 (N=9) and age matched CreER mice (N=9) was used for methylation clock assay. The methylation age of each mouse was normalized to its chronological age to calculate AgeAccel. p=0.0044, two-tailed Student’s t test. **d.** Frailty index scores of CreER and nmrHAS2 mice at 5- and 24-months of age. Pooled females and males. n=13-17. p-values were calculated with two-tailed Student’s t test. **e.** Rotarod performance of CreER and nmrHAS2 mice at 5- and 24-month of age. Pooled equal numbers of females and males, n=12. p-values were calculated with two-tailed Student’s t-test. **f.** Forelimb grip strength performance of CreER and nmrHAS2 mice at 5- and 24-month of age. Pooled females and males. n=13-17, p-values were calculated with two-tailed Student’s t-test. **g.** Old female nmrHAS2 mice have higher bone connectivity density. Hindlimb bones from 24 months old animals were taken for microCT scan. N=5-6. p-values were calculated with two-tailed Student’s t-test.

To further validate whether nmrHAS2 mice exhibit a younger biological age, we measured epigenetic age in livers of 24 months old mice using HorvathMammalMethylChip40^22, 23^. Epigenetic age was compared to chronological age to quantify age acceleration. Methylation age of CreER mice was close to their chronological age, while nmrHAS2 mice showed approximately -0.2 years of age acceleration (Fig. 2c, Extended Data Fig. 2c). The decreased age acceleration was observed in both males and females (Extended Data Fig. 2c-e). This result indicates that old nmrHAS2 mice have a significantly younger biological age than the chronological age. We also examined methylation levels of the 6553 CpG sites previously shown to undergo methylation changes during aging^24^. Our analysis revealed that, of the 6553 CpG sites, 165 were differentially methylated between nmrHAS2 mice and CreER mice. Among these sites, 145 sites that gain methylation during aging showed lower methylation in nmrHAS2 mice than in age-matched controls; and 20 CpG sites that lose methylation during aging showed higher methylation in nmrHAS2 mice than in age-matched controls (Extended Data Fig. 2f, g; Extended Data Table 2).

### nmrHAS2 mice have an improved healthspan

To provide a quantitative measure of health, we utilized a mouse frailty index (FI)^25^, which combined 31 parameters, including body weight, temperature, coat condition, grip strength, mobility, vision and hearing. The FI score increased with age for both nmrHAS2 and CreER mice. However, the FI score of old nmrHAS2 mice was substantially lower than that of the age-matched control group (Fig. 2d).

Rotarod performance^26^ was assessed to measure the locomotion and coordination of mice. The latency-to-fall time became significantly shorter for old CreER mice compared to the young CreER mice. However, old nmrHAS2 mice maintained youthful performance (Fig. 2e). We also measured the forelimb grip strength to evaluate the skeletal muscle function of the mice. The performance of both CreER and nmrHAS2 mice declined with age but old nmrHAS2 mice maintained a better performance compared to old CreER mice (Fig. 2f).

Osteoporosis is an important component of health span in females^27^. We performed micro-CT scan to assess the changes in the three-dimensional microstructure of the hindlimb of old mice. Connectivity density was shown to decrease during aging^28^. However, old nmrHAS2 female mice showed higher connectivity density in the femur, tibia, and tibia subchondral region (Fig. 2g). We did not observe the same phenotype in males (Extended Data Fig. 2h). Cumulatively, our results show that increased levels of HMW-HA in mice improves multiple parameters of healthspan.

### Gene expression analysis points to the reduced inflammation in nmrHAS2 mice

To investigate the mechanism responsible for increased lifespan and improved healthspan of nmrHAS2 mice, RNAseq analysis was performed on liver, muscle, white adipose tissue, kidney and spleen of 6- and 24-months old nmrHAS2 and control CreER animals (Fig. 3a). Expression patterns were compared across ages and genotypes (Fig. 3b, c). For most organs, nmrHAS2 mice showed fewer transcriptome changes during aging in both females and males which means the transcriptome of nmrHAS2 mice is less perturbed during aging (Fig. 3d).

**Figure 3.**
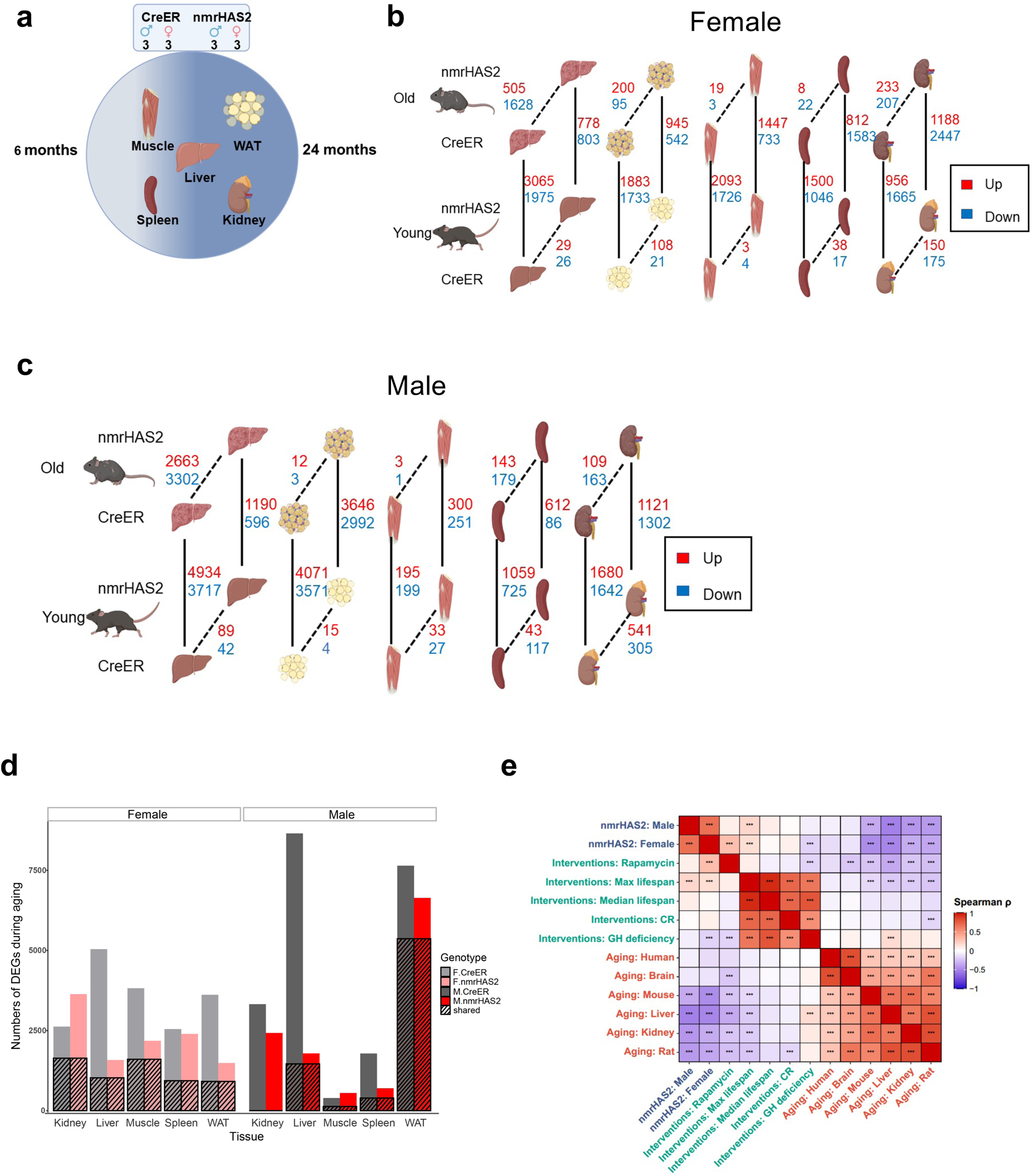
Transcriptome of nmrHAS2 mice undergoes fewer changes during aging than the transcriptome of CreER controls. **a.** Sequencing and sampling strategy. Liver, muscle, WAT, kidney and spleen 6- and 24-months old CreER and nmrHAS2 mice were used for RNA sequencing. Three biological replicates from each sex, age group, and genotype were used. **b.** Gene expression changes in females. Two parameters were compared: genotype, dashed lines; age, solid lines. **c.** Gene expression changes in males. Two parameters were compared: genotype, dashed lines; age, solid lines. **d.** Effects of aging on the transcriptome. The number of genes whose expression changed with age in either direction; hatched area represents genes that underwent changes in both genotypes. **e.** Association between nmrHAS2 effect on gene expression and signatures of lifespan-extending interventions and mammalian aging. Signatures of aging, lifespan-extending interventions and nmrHAS2 are shown in red, green and blue, respectively. *** p.adjusted < 0.001

We next asked if the transcriptome of nmrHAS2 mice shares any common features with transcriptomic changes induced by other pro-longevity interventions such as rapamycin, calorie restriction, and growth hormone knockout. We performed a hierarchical clustering analysis using RNAseq data from livers of young and old nmrHAS2 mice with expression data published for other interventions^29^. We bult a heatmap based on Pearson correlation coefficients across all datasets (Extended Data Fig. 3a). Interestingly, the transcriptomes of neither young nor old nmrHAS2 mice showed a clear correlation with any transcriptome data derived from the livers of mice subjected to other pro-longevity interventions. This result may suggest that increased levels of HMW-HA generated a novel pro-longevity transcriptomic signature (Extended Data Fig. 3a). To test whether the observed outcome is not due to noise generated by using the entire transcriptome data, we calculated Spearman correlation between top 400 gene expression changes induced by nmrHas2 expression and those associated with aging and established lifespan-extending interventions. We observed significant positive correlations between transcriptomic profiles of nmrHAS2 mice and signatures of rapamycin and murine maximum lifespan affected by longevity interventions (Fig. 3e). On the other hand, the effect of nmrHAS2 expression was negatively associated with multiple signatures of aging and biomarkers of interventions associated with growth hormone deficiency. The observed correlations were further amplified at the level of enriched pathways, estimated with Gene Set Enrichment Analysis (GSEA). Thus, at functional level nmrHAS2 signatures were positively associated with patterns of maximum and median lifespan, caloric restriction (CR) and rapamycin, and negatively correlated with all aging signatures (Extended Data Fig. 3b). Pro-longevity and anti-aging effects of nmrHAS2 expression were driven by significant downregulation of pathways associated with interleukin and interferon signaling, and by upregulation of genes involved in oxidative phosphorylation, respiratory electron transport and mitochondrial translation (Extended Data Fig. 3c, d). Remarkably, nmrHAS2 mice demonstrated stronger downregulation of inflammation and senescence than other examined lifespan-extending interventions. Taken together, our results suggest that nmrHAS2 expression in mice generates both pro-longevity and anti-aging transcriptomic changes, some of which are shared with other established interventions while others appear to be unique characteristics of nmrHAS2 model.

By reanalyzing the published transcriptome data from mice of different ages^30^, we determined the signature of gene expression changes during aging. The genes upregulated in old mice were defined as the ‘old’ gene set and the genes upregulated in young mice were defined as the ‘young’ gene set. Gene set enrichment analysis was conducted to compare old CreER and nmrHAS2 mice at the transcriptome level. Compared to old CreER mice that showed up-regulation of old gene sets in liver, old nmrHAS2 mice liver showed upregulation of young gene set (Fig. 4a). Although some tissues did not exhibit statistical significance, the trend displays high consistency across all tissues we sequenced (Extended Data Fig. 4a-e). This result indicates that tissues of old nmrHAS2 mice were shifted towards the ‘young’ state in both sexes at the transcriptomic level.

**Figure 4.**
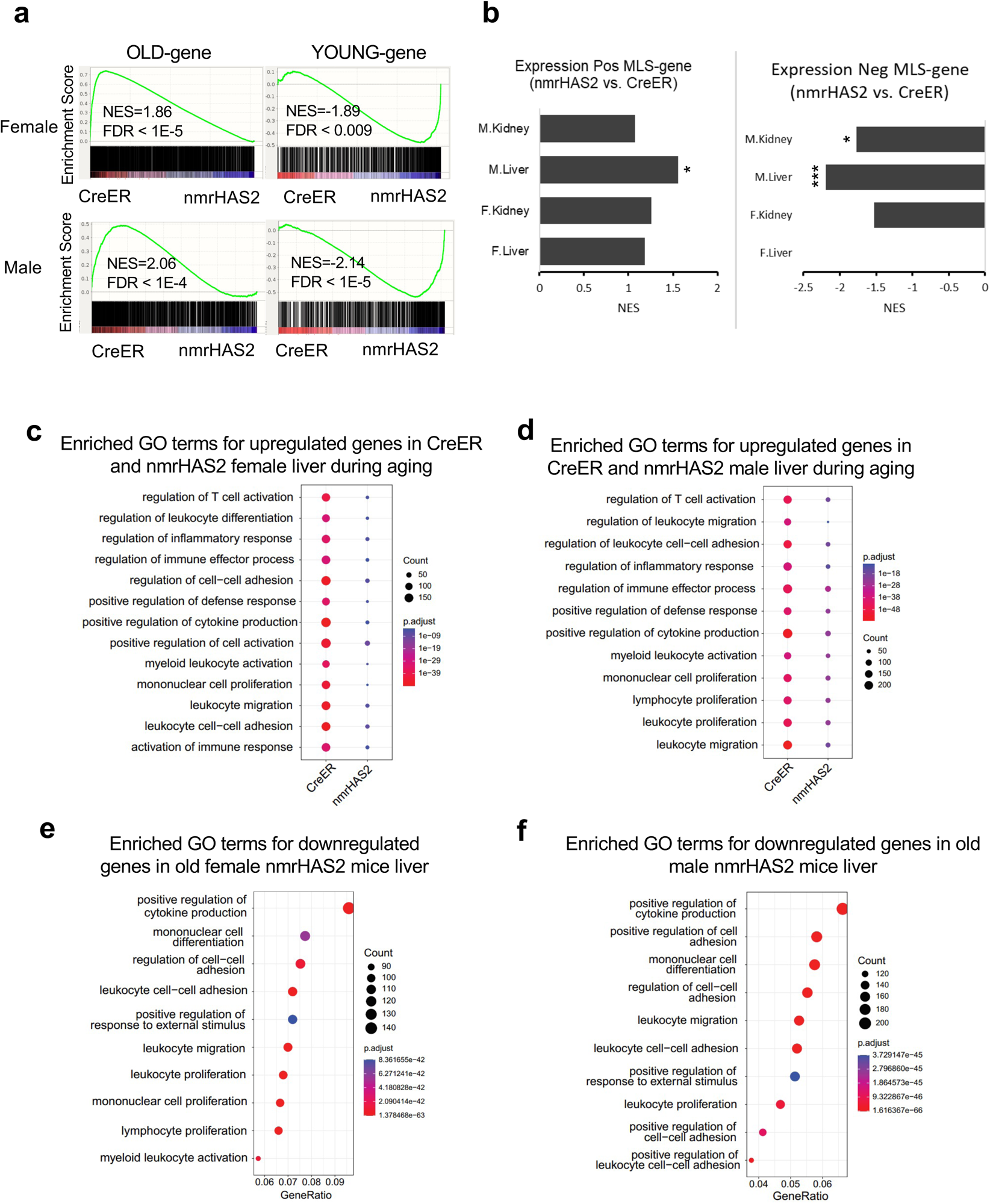
nmrHAS2 mice display younger transcriptome signature and reduced inflammation during aging. **a.** GSEA plots showing that YOUNG gene set is upregulated, and OLD gene set is downregulated in the liver of old nmrHAS2 mice of both sexes. **b.** GSEA shows that POS-MLS gene set was upregulated in liver and kidney of nmrHAS2 mice. NEG-MLS gene set is downregulated in liver and kidney of old nmrHAS2 mice. * FDR<0.05, *** FDR<0.001 **c.** GO term enrichment analysis shows that livers of old female nmrHAS2 mice have fewer inflammation-related pathways upregulated during aging. **d.** GO term enrichment analysis shows that livers of old male nmrHAS2 mice have fewer inflammation-related pathways upregulated during aging. **e.** GO term analysis shows that inflammation-related pathways are downregulated in livers of old female nmrHAS2 mice. **f.** GO term analysis shows inflammation-related pathways are downregulated in livers of old male nmrHAS2 mice.

In our previous comparative cross-species study, by analyzing the transcriptomes of 26 species, we obtained gene sets whose expression positively or negatively correlates with maximum lifespan named pos-MLS gene set and neg-MLS gene set, respectively^31^. By utilizing those gene sets for the GSEA, we found that the pos-MLS gene set is upregulated in livers and kidneys of old nmrHAS2 mice of both sexes. Conversely, the neg-MLS gene set was downregulated in the liver and kidneys of old male nmrHAS2 mice and in the kidneys of old female nmrHAS2 mice (Fig. 4b). This result suggests that nmrHAS2 transgene facilitates the expression of pro-longevity genes (genes highly expressed in long-lived species) and represses the expression of genes that are more highly expressed in short-lived species.

To further understand the mechanism of the anti-aging effects of hyaluronan, we compared genes whose expression changed with aging within each genotype. GO term enrichment analysis showed that CreER mice have more upregulated genes involved in the inflammatory response in the liver and spleen of both sexes, in kidneys for males, and in WAT and muscle for females (Fig. 4c, d, Extended Data Fig. 5a-h). This result indicates that HMW-HA attenuates age-related inflammation in multiple tissues.

**Figure 5.**
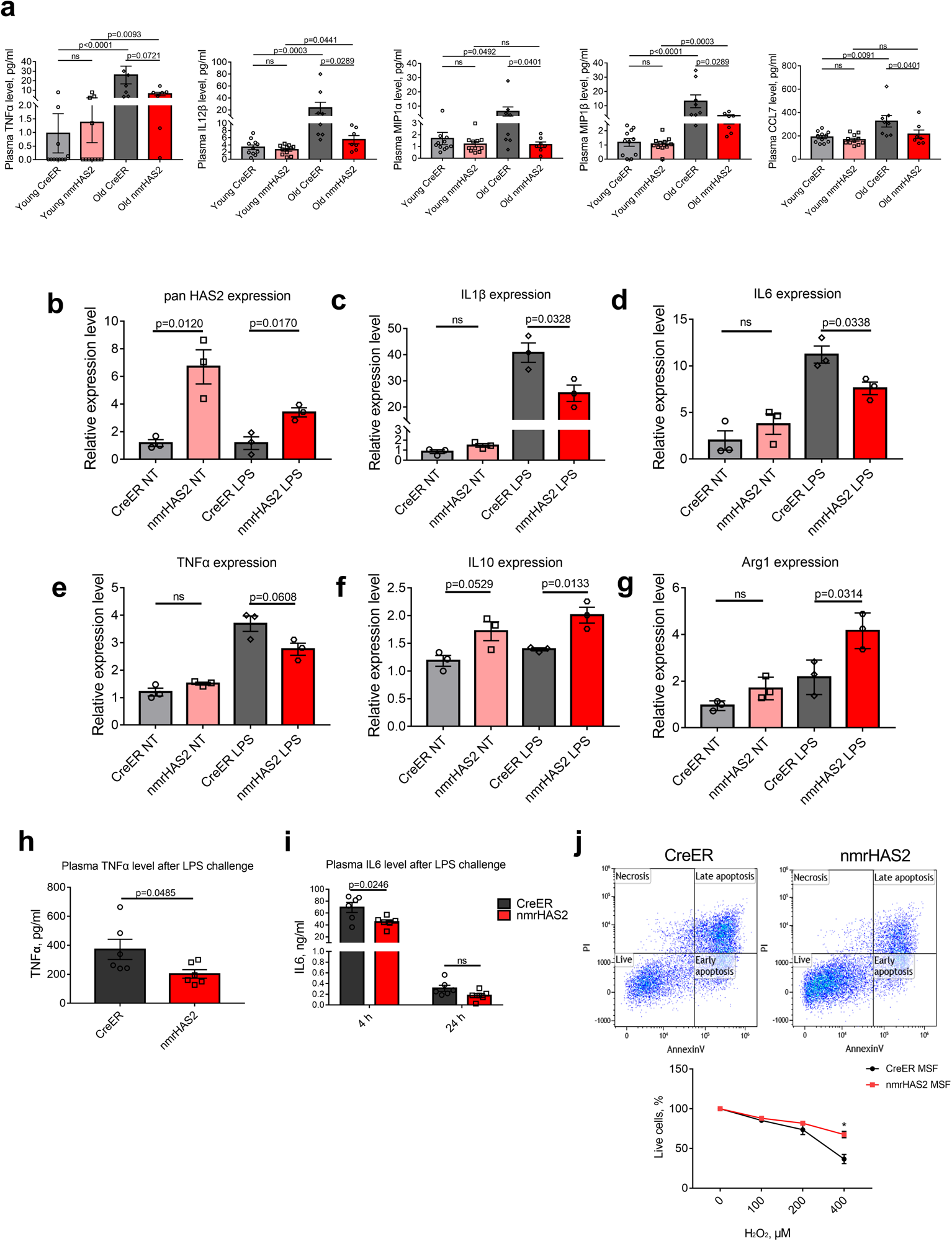
nmrHAS2 reduces pro-inflammatory response in *vitro* and in *vivo* and protects cells from oxidative stress. **a.** Luminex multiplex immunoassay shows that old female nmrHAS2 mice have reduced levels of multiple inflammatory cytokines and chemokines. N=7-11. p-values were calculated using Mann-Whitney u-test. **b.** BMDM from nmrHAS2 mice have significantly upregulated HAS2 levels. BMDMs were isolated from 5-months old male mice (N=3). p-values were calculated using two-tailed Student’s t-test. **c.** BMDM from nmrHAS2 mice have lower levels proinflammatory IL-1β mRNA after LPS challenge. BMDMs were isolated from 5-months old male mice (N=3). p-values were calculated using two-tailed Student’s t-test. **d.** BMDM from nmrHAS2 mice have lower levels of proinflammatory IL6 mRNA after LPS challenge. BMDMs were isolated from 5-months old male mice (N=3). p-values were calculated using two-tailed Student’s t-test. **e.** BMDM from nmrHAS2 mice have higher levels of proinflammatory TNFα mRNA after LPS challenge. BMDMs were solated from 5-months old male mice (N=3). p-values were calculated using two-tailed Student’s t-test. **f.** BMDM from nmrHAS2 mice have higher levels of anti-inflammatory IL10 mRNA after LPS challenge. BMDMs were isolated from 5-months old male mice (N=3). p-values were calculated using two-tailed Student’s t-test. **g.** BMDM from nmrHAS2 mice have higher levels of anti-inflammatory Arginase 1 mRNA after LPS challenge. BMDMs were isolated from 5-months old male mice (N=3). p-values were calculated using two-tailed Student’s t-test. **h.** nmrHAS2 mice have significantly lower plasma TNFα level 4 h after LPS challenge. 5-months old male mice (N=3). p-values were calculated using two-tailed Student’s t test. **i.** nmrHAS2 mice have significantly lower plasma IL6 levels 4 h after LPS challenge. 5-months old male mice (N=3). p-values were calculated using two-tailed Student’s t-test. **j.** Skin fibroblast cells isolated from nmrHAS2 mice are more resistant to H_2_O_2_ treatment. Skin fibroblasts were isolated from 5-months old male mice. N=3. p-values were calculated using two-tailed Student’s t-test.

Additionally, we analyzed the differentially expressed genes (DEGs) between two genotypes of mice at the same age. Expression of nmrHAS2 had very mild effects on the overall transcriptome of young mice, and there were very few differentially DEGs observed between young nmrHAS2 and CreER mice (Fig. 3b, c). For old mouse organs, we picked the liver which showed most DEGs between nmrHAS2 and CreER mice for the GO term enrichment analysis. Our results revealed that both female and male nmrHAS2 liver showed reduced expression of inflammatory-related genes and higher expression of genes involved in normal liver functions such as nutrient metabolism. This result indicates that the liver of old nmrHAS2 mice showed reduced inflammation and better-preserved functions compared to controls (Fig. 4e, f; Extended Data Fig.5 i, j). Overall, these results demonstrate that nmrHAS2 mice display reduced age-related inflammation.

### HA reduces the inflammatory response *in vitro* and *in vivo* and protects cells from oxidative stress

To validate whether nmrHAS2 mice show reduced inflammation during aging, we collected plasma from young and old mice and performed Luminex multiplexing cytokine assay to test thirty-six cytokines and chemokines. Several targets showed higher levels in males, but the trend did not reach statistical significance due to high individual variability (Extended Data Fig. 6). In females, almost all cytokine and chemokine levels were increased during aging which is consistent with the effect of sex hormones on immunity during aging^32^. Remarkably, the majority of pro-inflammatory cytokines and chemokines were lower in old nmrHAS2 mice compared to the age-matched controls (Extended Data Fig. 7). The differences for pro-inflammatory cytokines IL12p40, MIP1*α*, MIP1*β*, and chemokine CCL7 reached statistical significance (Fig. 5a). Collectively, the transcriptome and cytokine data show that overexpression of nmrHAS2 attenuates inflammaging in mice.

**Figure 6.**
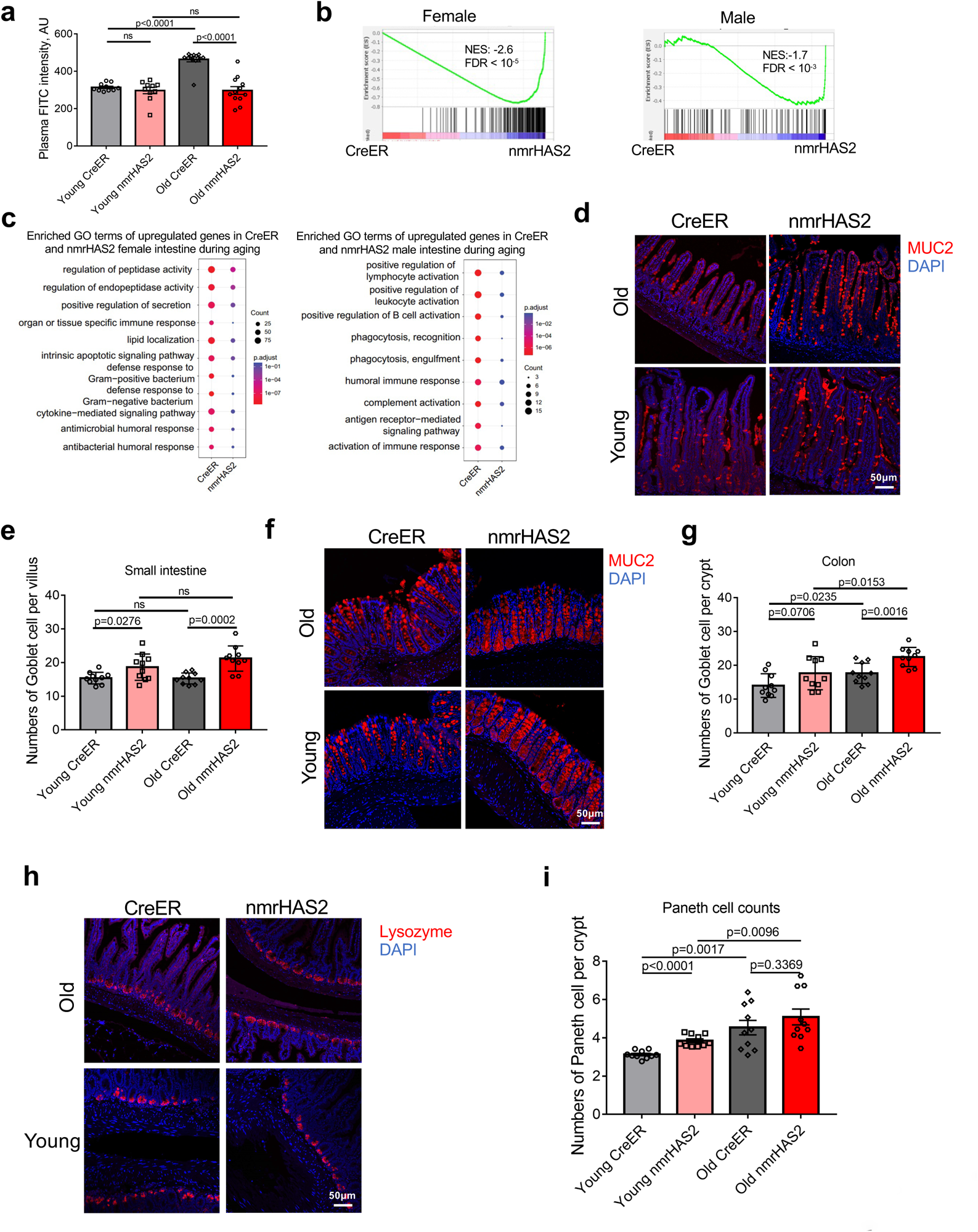
nmrHAS2 mice are protected from age-related loss of gut barrier function. **a.** nmrHAS2 mice have less leaky gut compared to the age matched controls. Pooled females and males (N=10-12). p-values were calculated using two-tailed Student’s t-test. **b.** GSEA shows that old nmrHAS2 mice have a younger intestine at the transcriptome level for both sexes. **c.** GO term analysis shows that small intestine of old nmrHAS2 mice has fewer inflammatory-related pathways upregulated during aging for both sexes. **d.** Representative pictures of Goblet cell staining in the small intestine of nmrHAS2 and CreER mice. **e.** Quantification of goblet cells in the small intestine of 7- and 24-months old mice (shown in D). Pooled females and males (N=10). p-values were calculated using two-tailed Student’s t-test. **f.** Representative pictures of Goblet cell staining in the distal colon of nmrHAS2 and CreER mice. **g.** Goblet cell counts in the distal colon of 7- and 24-month-old mice (shown in F). Pooled females and males (N=10). p-values were calculated using two-tailed Student’s t-test. **h.** Representative pictures of Paneth cell staining in the small intestine of nmrHAS2 and CreER mice. **i.** Paneth cells counts in the small intestine of 7- and 24-months old mice. Pooled females and males (N=10). p-values were calculated using two-tailed Student’s t-test.

**Figure 7.**
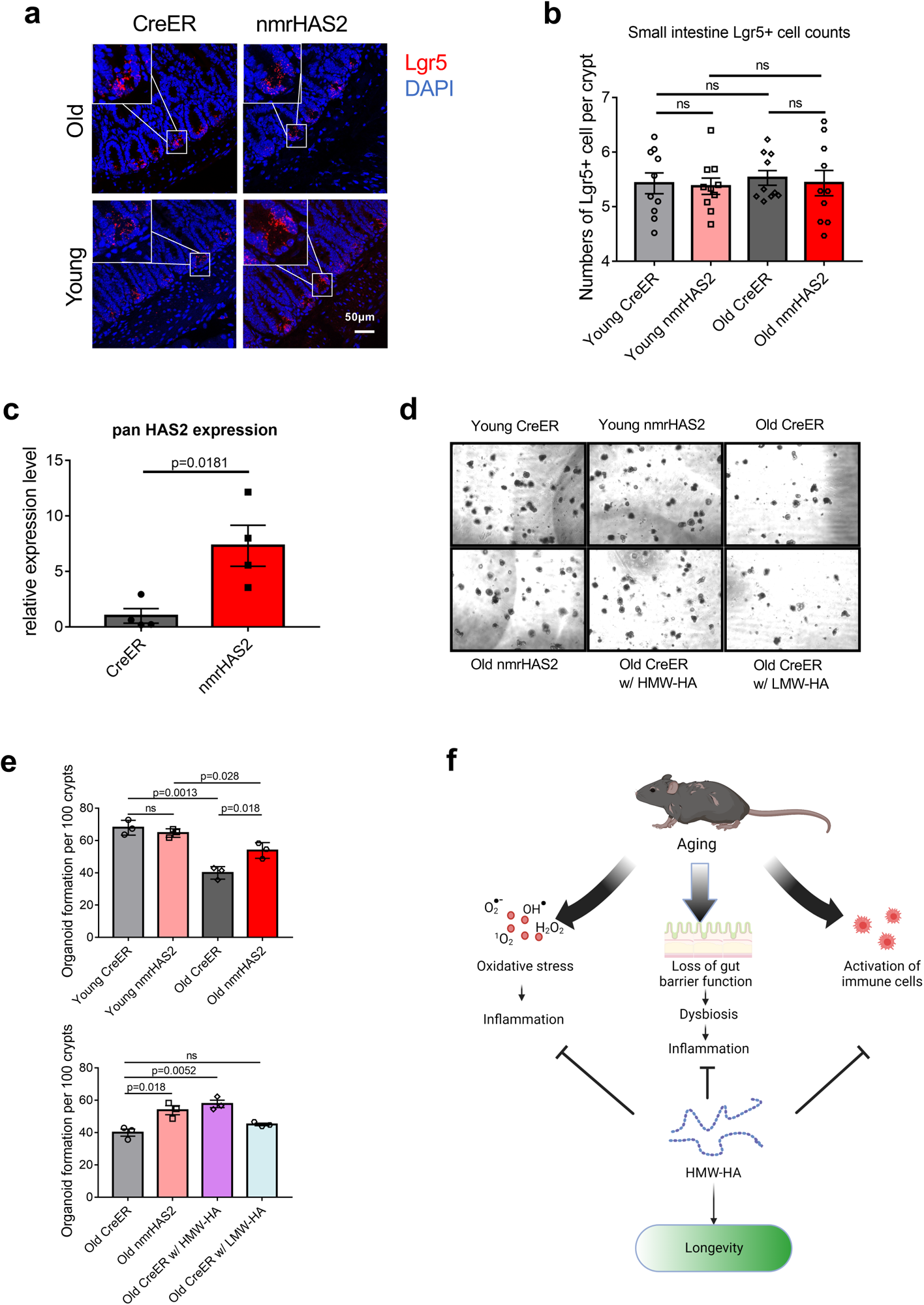
HMW-HA improves maintenance of intestinal stem cells during aging. **a.** Representative pictures of Lgr5 *in situ* hybridization in the small intestine of young and old nmrHAS2 and CreER mice. **b.** Lgr5+ intestinal stem cell counts in the small intestines of 7- and 24-months old mice. Pooled females and males (N=10). P-values were calculated using two-tailed Student’s t-test. **c.** Intestinal crypts isolated from nmrHAS2 mice have significantly upregulated HAS2 levels. Intestinal crypts were isolated from 5-months old mice (N=4). p-values were calculated using two-tailed Student’s t-test. **d.** Intestinal crypts from old nmrHAS2 mice form more intestinal organoids *in vitro* N=3. Addition of HMW-HA, but not LMW-HA, to CreER crypts resulted in a greater number of organoids. **e.** Organoid quantification in 7- and 24-months old mice (N=3). p-values were calculated using two-tailed Student’s t-test. **f.** Model for anti-aging effects of HMW-HA. HMW-HA produced by overexpression of nmrHAS2 gene protects cells from oxidative stress, improves maintenance of intestinal stem cells to provide a better gut barrier function during aging, and reduces the production of pro-inflammatory molecules by immune cells. The beneficial effects of HMW-HA further contribute to longevity and healthspan of the mice.

HMW-HA molecules exert anti-inflammatory and immunoregulatory effects^33^. It was reported that HMW-HA represses classic pro-inflammatory M1 macrophage activation but promotes an anti-inflammatory alternative M2 macrophage activation^34^. We hypothesized that macrophages isolated from nmrHAS2 mice may have reduced response to the pro-inflammatory stimuli. To test this hypothesis, we isolated bone marrow-derived macrophages (BMDM) from young CreER and nmrHAS2 mice and cultured them in *vitro*. Macrophages from male nmrHAS2 mice showed a 6-folds increase in panHAS2 mRNA expression compared to macrophages from male CreER mice (Fig. 5b). To check the activation of macrophages, we treated bone marrow-derived macrophages with *E. coli* lipopolysaccharide (LPS). The HAS2 level dropped after LPS treatment but remained significantly higher for nmrHAS2 macrophages isolated from male mice. Two major HAases HYAL1 and HYAL2 level also dropped after LPS treatment (Extended data Fig. 8c-f). nmrHAS2 macrophages from males produced significantly lower levels of pro-inflammatory IL1*β* and IL6 (Fig. 5c, d) compared to the CreER macrophages. TNF*α* also showed lower levels in nmrHAS2 cells but the effect did not reach statistical significance (Fig. 5e). Interestingly, male nmrHAS2 macrophages produced higher levels of anti-inflammatory IL10 and arginase 1 after LPS challenge (Fig. 5f, g). To test whether the anti-inflammatory effect is due to increased HMW-HA rather than to an unknown function of nmrHAS2, we generated Raw264.7 macrophage cell lines overexpressing either mouse HAS2 or nmrHAS2 under the control of the same CAG promoter and challenge them with LPS. Macrophages overexpressing any form of HAS2 exhibited an anti-inflammatory impact similar to that seen in primary macrophages, implying that the anti-inflammatory effect arose from the production of high molecular weight hyaluronic acid (HMW-HA) (Extended data Fig. 8b).

Macrophages from female nmrHAS2 mice had a lower HAS2 level compared to macrophages from nmrHAS2 males. After treatment with LPS, nmrHAS2 levels in female nmrHAS2 macrophages dropped to the same level as that in female CreER macrophages (Extended data Fig. 8a). Consequently, females did not show expression differences in cytokines after LPS challenge (data not shown). LPS stimulation triggered similar HAS2 and hyaluronidase (HAase) changes in both BMDMs and macrophage cell lines (Extended data Fig. 8g-i). The elevation of HA levels upon LPS challenge could be attributed to the decline in hyaluronidase (HYAL) expression. Moreover, the LPS-treated media derived from HAS2-overexpressing macrophages exhibited a substantial accumulation of HMW-HA (Extended data Fig. 8j, k). Consequently, the accumulation of HMW-HA produced by HAS2 during macrophage activation likely accounts for the anti-inflammatory effects.

To investigate whether HMW-HA confers an anti-inflammatory effect *in vivo*, we examined the response of nmrHAS2 and CreER mice to the LPS. When young mice were injected i.p. with low dose LPS, male mice had a stronger response compared to females as evidenced by the higher level of plasma TNF*α* 4 hours after the treatment (Fig. 5h, Extended data Fig. 9a). Consistent with the *in vitro* results, both male and female nmrHAS2 mice showed reduced plasma TNF*α* levels 4 hours after LPS injection (Fig. 5h, Extended data Fig. 9a). Female nmrHAS2 mice had lower levels of plasma TNF*α* 24 hours after the injection (Extended data Fig. 9a). In addition, both male and female nmrHAS2 mice produced less IL6 in plasma 4 h after LPS treatment (Fig. 5i, Extended data Fig. 9b). We observed reduced inflammation in multiple tissues of female nmrHAS2 mice 24 hours after injection. Liver, spleen, and kidney from female nmrHAS2 mice showed significantly lower pro-inflammatory IL1*β*, TNF*α*, but similar IL6 mRNA levels (Extended data Fig. 9c-e). These results indicate that HMW-HA suppresses pro-inflammatory response of nmrHAS2 mice both *in vitro* and *in vivo* which contributes to reduced inflammation in old nmrHAS2 mice.

HMW-HA protects cells from oxidative stress^13^. Primary fibroblasts from nmrHAS2 mice produced more abundant hyaluronan (Extended data Fig. 9f, g). Consistently, nmrHAS2 cells showed higher survival after H_2_O_2_ treatment indicating the protective effect of HMW-HA against oxidative stress (Fig. 5j, Extended data Fig. 9h). To test that the protective effect is conferred by HMW-HA and not by another function of nmrHAS2 gene, we generated a mouse HAS2 overexpressing fibroblast cell line. Overexpression of the mouse HAS2 also resulted in the increased production HMW-HA as nmrHAS2 and exhibited a similar protective effect against oxidative stress (Extended data Fig. 10a-c). As oxidative stress is linked to inflammation, we hypothesize that an additional pathway by which HMW-HA counteracts inflammation is by reducing oxidative stress.

### HMW-HA improves intestinal stem cell maintenance and preserves intestinal health during aging

Disruption of gut barrier in old individuals contributes to chronic inflammation during aging and promotes age-related diseases^35, 36^. Since old nmrHAS2 mice showed reduced inflammation in multiple organs, we compared the gut barrier function of nmrHAS2 and control mice. The gut permeability increased with age in CreER mice as measured by the gut to blood transfer of FITC-dextran signal. Remarkably, gut permeability remained unchanged between old and young nmrHAS2 mice (Fig. 6a). Gene set enrichment analysis of the transcriptome of the small intestine from young and old mice showed that nmrHAS2 mice have the transcriptome shifted towards the younger state (Fig. 6b). GO term analysis showed reduced inflammation during aging in nmrHAS2 mice of both sexes (Fig. 6c).

The intestinal epithelium contributes to the maintenance of intestinal barrier function. Loss of functional epithelial cells leads to the leaky gut^37^. The mucus layer secreted by goblet cells provides a physical barrier preventing interactions between gut bacteria and intestinal epithelial cells. Remarkably, both young and old nmrHAS2 mice had more goblet cells in their small intestine and colon compared to the age-matched CreER controls (Fig. 6d-g). Paneth cells secrete antibacterial peptides, which provide a chemical barrier in the small intestine^38^. Paneth cell number increased with age in both nmrHAS2 and CreER mice, which is believed to be an adaptive response to age-associated gut dysbiosis^39^. Interestingly, young nmrHAS2 mice had more Paneth cells than the young control mice and displayed a lower relative increase in the Paneth cell number with age (Fig. 6h, i). The smaller age-related increase in the Paneth cell number in nmrHAS2 mice may be due to improved intestinal health.

Intestinal stem cells (ISCs) located in the crypts give rise to the goblet, Paneth cells, and absorptive enterocytes. Loss of functional ISCs during aging is an important contributor to age-associated gut dysbiosis^40^. nmrHAS2 and control mice had similar numbers of ISCs in young age and old age (Fig. 7a, b). However, we observed higher expression of WNT and Notch pathway related genes in old nmrHAS2 mice intestine suggesting better stem cell maintenance (Extended data Fig. 11a). To test our hypothesis, we then tested the functionality of the ISCs by their ability to form organoids. Crypts from old CreER mice formed much fewer organoids than the crypts from young CreER and nmrHAS2 mice. Remarkably, the crypts isolated from nmrHAS2 mice showed a strong expression of HAS2 gene (Fig. 7c), and the ability of those crypts to form organoids did not decline with age (Fig. 7d, e). To test whether the improved stemness of ISCs in nmrHAS2 mice was due to hyaluronan, we added HMW-HA or LMW-HA into the Matrigel used to culture organoids. Supplying HMW-HA was sufficient to reactivate ISCs from old CreER mice and resulted in a higher number of organoids (Fig. 7e, f). This result indicates that HMW-HA produced by nmrHAS2 helps maintain stemness of ISCs during aging.

The gut microbiome undergoes changes during aging and gut dysbiosis can further contribute to systemic inflammation^41^. Since the old nmrHAS2 mice showed a heathier gut, we compared the microbial composition between 7- and 24-months old nmrHAS2 and control mice by sequencing 16S rDNA. At the phylum level, the Operational Taxonomic Units (OTUs) showed that both young and old nmrHAS2 mice had increased *Bacteroidetes* and decreased *Firmicutes* levels compared to age-matched controls. However, only the old groups reached statistical significance (Extended data Fig. 11b, c). Decrease of *Bacteroidetes* to *Firmicutes* ratio was shown to correlate with gut dysbiosis in hypertension and metabolic disorders^42, 43^. At the family level, *Deferribacteraceae*, *Streptococcaceae*, and *Lachnospiraceae,* which are known to positively correlate with inflammation, showed higher abundance in old CreER mice. *Muribaculaceae* which was linked to longevity of *Spalax leucodon*^44^, was found at higher levels in old nmrHAS2 mice (Extended data Fig. 11d, e). Collectively, our results indicate that old nmrHAS2 mice have improved intestinal health, contributing to reduced age-related inflammation.

## Discussion

In the tissues of the naked mole-rat, HMW-HA is abundant and contributes to cancer resistance and possibly longevity of this exceptionally long-lived rodent^1, 2^. Here we demonstrated that this evolutionary adaptation, unique to the naked mole-rat, can be “exported” to other species. The HMW-HA produced by naked mole-rat HAS2 gene conferred cancer resistance and increased lifespan of mice. nmrHAS2 mice accumulated HMW-HA in multiple major organs; these mice were resistant to spontaneous and chemically induced cancer and showed an extended median and maximum lifespan. nmrHAS2 mice also displayed improved healthspan including lower frailty scores and physical performance. This was accompanied by lower methylation age. The transcriptomes of aged nmrHAS mice had more youthful features when compared to the aging signature derived from the Tabula Muris Senis data^30^. The change conferred by nmrHAS2 was distinct from the transcriptomic changes triggered by known life-extending interventions including rapamycin, acarbose, growth hormone deletion, calorie restrictions and others. Interestingly, the nmrHAS2 altered the mouse transcriptome in a direction consistent with the transcriptomic signature of the long-lived species^31^. We therefore hypothesize that the presence of HMW-HA changes the transcriptome towards that of a longer-lived species (Fig. 4a).

What is the molecular mechanism by which HMW-HA extends mouse lifespan and healthspan? Importantly, due to the high conservation of the HAS2 gene, we do not believe that the naked mole-rat sequence *per se* was critical for the pro-longevity effect of nmrHAS2. Expressing mouse HAS2 gene showed the same protective effects *in vitro*. Rather, the increased production of HMW-HA was important. The most striking difference we observed between the transcriptomes of the aged nmrHAS2 mice and their littermate controls was the downregulation of multiple pathways related to inflammation in most tissues we analyzed. Beneficial effects of HMW-HA have recently been reported for muscle stem cells and adipose tissues where HMW-HA reduced inflammatory signaling and improved glucose homeostasis, respectively^45, 46^. Interestingly, some of the effects were systemic in nature suggesting that even the tissues not displaying elevated HA levels can benefit from elevated HA elsewhere in the body^45, 46^. Additionally, the plasma of aged nmrHAS2 mice also displayed reduced levels of multiple pro-inflammatory cytokines which indicates the systemic anti-inflammatory effect of HMW-HA. Age-associated chronic inflammation, so-called inflammaging, contributes to the pathogenesis of age-related diseases^47, 48^. It has been reported that individuals which have higher than age-average levels of inflammatory markers are more fragile and likely to be hospitalized^49^.

The lifespan extension we observed in nmrHAS2 mice was significant but modest, while the healthspan improvement was more robust. While we observed strong expression of nmrHAS2 across all tissues, HA accumulation was modest and did not reach the levels observed in the naked mole-rat. This is likely explained by the high activity of hyaluronidases in mouse tissues^2, 50^. In the naked mole-rat high level of HA is achieved by both robust synthesis and very slow degradation^2^, yet in our model only the synthesis arm was modified. We hypothesize that if we were able to simultaneously attenuate HA degradation in nmrHAS2 mice we would achieve a greater lifespan extension.

We then discovered two independent pathways by which HMW-HA conferred an anti-inflammatory effect to the mice. First, HMW-HA had a direct immunomodulating effect on the immune cells of nmrHAS2 mice. Monocyte–macrophage lineage cells act as major effector cells in chronic inflammatory processes in aging-related diseases^51^. Macrophages are often divided into two subgroups: M1 and M2^52^. Aging may modulate M1/M2 activation and polarization. Sustained activation of M1 macrophage is linked to tissue dysfunction, while M2 macrophage promotes tissue homeostasis^53^. HMW-HA has been shown to prime macrophages towards the M2 state^34, 54, 55^. We also observed the same phenotype when we primed the bone marrow-derived macrophages from nmrHAS2 mice with *E. coli* lipopolysaccharide. BMDMs from nmrHAS2 mice showed a much higher expression of HAS2 gene, which may potentially help them to produce more HMW-HA and facilitates the alternative activation. HMW-HA was also shown to play an immunoregulatory role *in vivo* by binding to several immune cell types^56^. For instance, HMW-HA promotes the expression of FoxP3 in Treg cells by crosslinking CD44 which helps to maintain immunologic tolerance^57^. HMW-HA binds to CD44 and hSiglec-9 receptor which suppresses neutrophil extracellular trap formation and oxidative burst^58^. We observed that nmrHAS2 mice produced a lower level of inflammation after the LPS challenge which is consistent with previous reports that intraperitoneal injection of HMW-HA protects mice from LPS-induced sepsis^14^.

The second, unexpected pathway, by which HMW-HA conferred its anti-inflammatory effect was through improved intestinal health. Aged nmrHAS2 mice were protected from leaky gut, and had healthier microbiomes. One of the major contributors to inflammaging is believed to be age-related deterioration of the intestinal barrier and translocation of gut bacterial products to the bloodstream triggering an immune response^36^. Intestinal barrier failure is associated with aging-related systemic disorders such as obesity, metabolic disorder, brain dysfunction, and cancer^35, 59^. Reduced thickness of the mucus layers was considered to be the cause of leaky gut^60^. It has been shown that the colonic mucus layer decreased in aging mice, suggesting an association with bacterial penetration and immune activation^61, 62^. Interestingly, we found that elevated production of hyaluronan increased the number of mucin-producing goblet cells in both small intestine and colon, suggesting old nmrHAS2 are protected from leaky gut due to increased mucin formation. The increase in goblet cells in nmrHAS2 mice was significant, but very mild, and did not resemble a pathological over-proliferation of goblet cells observed in a disease state such as cystic fibrosis^63^.

Studies in *Drosophila* showed that with aging intestinal stem cell (ISC) lose their stemness, are no longer able to differentiate into functional intestinal cells and undergo hyperproliferation associated with a loss of barrier function^64–66^. While the hyperproliferation of ISCs has not been unequivocally demonstrated in mammals^64, 67^, in mammals aging may be similarly associated with the loss of stemness by the ISCs. Indeed, it was found that the regenerative capacity of ISCs from old mice^67^ and old human^64^ was diminished *in vitro*. Consistent with these prior reports, we observed that ISCs from aged wild type mice formed fewer intestinal organoids. Remarkably, there was no such decrease observed for nmrHAS2 mice. Interestingly, the decline in the formation of organoids in the wild type mice could be rescued by adding HMW-HA to the culture media, indicating that the presence of HMW-HA promotes stemness of aged ISCs. It was reported that intraperitoneal injection of HA to mice promotes Lgr5+ stem cell proliferation and crypt fission through CD44 and TLR4^68, 69^. Collectively, our results suggest that HMW-HA produced by nmrHAS2 transgene improves the maintenance of ISCs resulting in a healthier gut barrier during aging. Consequently, the healthier gut barrier function inhibits the shift of the gut microbiome towards proinflammatory commensals and reduction of beneficial microbes, slowing down the onset of inflammaging.

The resistance of nmrHAS2 mice to both spontaneous and induced cancer may be driven by both cell-autonomous mechanisms resulting from anti-proliferative signaling of HMW-HA through the CD44 receptor^70^, and by its systemic anti-inflammatory effect^12^. Indeed, it was reported that HMW-HA blocks melanoma cell proliferation by signaling through CD44 to promote G1/G0 arrest^71^. HMW-HA also suppresses the growth of murine astrocytoma cell lines, glioma and colon carcinoma xenografts^72^. HMW-HA has also been reported to reduce the migratory and invasive capacity of aggressive cancer cells^73, 74^. On the other hand, chronic inflammation plays an important role in the development of cancer^47^. Many studies have shown that inflammatory cells can promote the occurrence and development of tumors by facilitating cancer cell proliferation, angiogenesis, and tumor invasion^75–77^. The anti-inflammatory properties of HMW-HA discussed above may potentially reduce the chronic inflammation during aging and prevent cancer initiation. Additional pro-longevity effects of HMW-HA can be linked to its antioxidant and cytoprotective properties. Indeed, HMW-HA was reported to enhance cellular oxidative stress resistance^78, 79^. Consistent with these observation fibroblasts from nmrHAS2 mice showed resistance to oxidative stress.

In summary, our results demonstrate that HMW-HA produced by the nmrHAS2 gene extends lifespan and improves healthspan of mice by ameliorating age-related inflammation. This is achieved by directly suppressing the production of pro-inflammatory factors by immune cells, and by promoting stemness of ISCs preventing age-related decline of the intestinal barrier (Fig. 7f). These findings demonstrate that evolutionary adaptations found in long-lived species such as the naked mole-rat can be exported and adapted to benefit human health. Additionally, these findings underscore the utility of HMW-HA for treatment of age-related inflammation in the intestine and other tissues.

## Acknowledgements

This work was supported by grants from the National Institutes of Health to V.N.G., A.S. and V.G.

## Author contributions

Z.Z., A.S., and V.G. designed research, analyzed data, and wrote the manuscript; Z.Z. performed most of the experiments. Y.L., A.T., V.N.G. analyzed RNAseq data. X.T. designed research, generated the transgenic mouse strain. Z.Z. X.T. F.T. and S.E. performed the aging study. K.B. helped with immunofluorescent staining. J.A. performed the DMBA/TPA treatment. D. F. helped with cell apoptosis assay. E.R., S.B. helped with maintaining the mouse colony. S. H. performed the methylation clock assay. A.S. and V.G. supervised research.

## Methods

### Animal husbandry

All animal experiments were approved and performed in accordance with guidelines set forth by the University of Rochester Committee on Animal Resources with protocol number 2017-033 (mouse). Mice were group housed in IVC cages (up to 5 animals/cage) in a specific pathogen-free environment and fed standard chow diet (Altromin 1324; total pathogen free, irradiated with 25 kGy) and water ad libitum. Animal rooms were maintained at 21–24 °C and 35–75% relative humidity, with 12/12 h (6 a.m. to 6 p.m.) dark–light cycle. Cages were routinely replaced every 10–14 days.

### Mice and lifespan study

C57BL/6 mice were from Charles River Labs, R26-CreERT2 mice were from the JAX. To generate nmrHAS2 conditional transgenic mice, nmrHAS2 coding sequence was subcloned into the pCALNL-GFP plasmid (Addgene plasmid # 13770) to replace GFP. Transgenic mice were made by UC-Irvine Transgenic Mouse Facility. nmrHAS2 and control CreER mice were obtained by crossing mice heterozygous for nmrHAS2 gene with homozygous R26-CreERT2 mice. At the age of 1 month, progenies were separated by sex, ear tagged, and distal tail (∼2 mm) was cut for genotyping determination. All mice received 80mg/kg tamoxifen at 3 months of age for 5 consecutive days. CreER and nmrHAS2 mice were housed in the same cage for all experiments. None of the animals entered into the aging study were allowed to breed. Mice were inspected daily for health issues, and any death was recorded. Animals showing significant signs of morbidity, based on the AAALAC guidelines, were euthanized for humane reasons and were used for lifespan analysis since they were deemed to live to their full lifespan. No mice were censored from analysis. Lifespan was analyzed by Kaplan–Meier survival curves, and p-values were calculated by log-rank test using Graphpad prism.

### Necropsy

Cages were inspected every night. Dead animals were removed from cages, opened, and examined macroscopically by a trained person. A fraction of animals could not be examined because they were too decomposed or disturbed by other animals. Organs were moved, turned, or lifted with forceps for the examination but were not removed. All visible tumors, as well as any other observations were noted.

### Tissue and plasma collection

Animals were brought to the laboratory in their holding cages and euthanized one by one for dissection. Mice were euthanized by isofluorane anesthesia followed by cervical dislocation. The dissection was performed as rapidly as possible following euthanasia by several trained staff members working in concert on one mouse. Tissue samples were either rapidly frozen in liquid nitrogen (for HA amount and MW determination, and RNA sequencing) or fixed in 4% formalin (for histology). Blood was collected by cardiac puncture into EDTA-coated tubes, centrifuged, and the plasma was aliquoted and rapidly frozen in liquid nitrogen. All frozen samples were stored at -80*℃*.

### HA preparation

For purifying HA from tissues, 200mg of pulverized tissue from 5 months old mice were mixed with proteinase K solution (final concentrations of 1 mM Tris-Cl pH 8.0, 2.5 mM EDTA, 10 mM NaCl, 0.05% SDS, 2 mg/ml proteinase K), and incubated at 55 °C overnight followed by saturated phenol-chloroform-isoamyl alcohol (Sigma) extraction. HA was precipitated with isopropanol and centrifugation (12,000 × g for 15 min) then wash with 70% ethanol (12,000g 10min) and dissolved in 600ul 10mM Tris buffer at PH 8 overnight at RT. The purified HA was digested with SuperNuclease (final concentration of 50U/ml, Lucerna-chem) overnight at 37*℃* to eliminate nucleic acid contamination. HA was extracted by saturated phenol-chloroform-isoamyl alcohol (Sigma), precipitated by isopropanol, and washed with 70% ethanol again. The pellet was dissolved in 30ul 10mM Tris buffer at PH 8 overnight at RT.

For HA purification from media, conditioned media were first mixed with proteinase K solution (final concentrations of 1 mM Tris-Cl pH 8.0, 2.5 mM EDTA, 10 mM NaCl, 0.05% SDS, 1 mg/ml proteinase K) and incubated at 55 °C for 4 h. Following protein digestion, media were extracted with saturated phenol-chloroform-isoamyl alcohol (Sigma). HA was precipitated with ethanol and centrifugation (4,000 × g for 45 min). HA pellet was dissolved in PBS and then extracted with 1/100 volume of Triton-X114. After Triton-X114 extraction, HA was precipitated again with ethanol. Finally, HA pellet was washed with 70% ethanol and dissolved in PBS.

### Pulse-field gel electrophoresis

Purified HA was mixed with sucrose solution (final concentration of 333 mM) and loaded to a 0.4% SeaKem Gold agarose gel (Lonza). HA-Ladders (Hyalose) were run alongside the samples. Samples were run 16 h at 9 °C at 4 V with a 1–10 running ratio in TBE buffer using CHEF-DRII system (Bio-Rad). After the run, the gel was stained with 0.005% (w/v) Stains-All (Santa Cruz) in 50% ethanol overnight. Then the gel was washed twice with 10% ethanol for 12h, exposed to light to decrease background, and photographed with ChemiDoc Imaging System (Bio-Rad).

### HA ELISA

Hyaluronan concentration in the media was quantified using the hyaluronan ELISA kit (R&D systems) following the manufacturer’s instructions.

### Rotarod performance

Motor performance was assessed using the protocol described before^80^. Briefly, gross motor control was measured using the rotarod (IITC Life Science, CA, USA). For this test, each mouse was placed on a cylindrical dowel (95.525 mm in diameter) raised ∼30 cm above the floor of a landing platform. Mice were placed on the dowels for 3 min to allow them to acclimatize to the test apparatus. Once initiated the cylindrical dowels began rotating and accelerated from 5 rpm to a final speed of 20 rpm over 60 s. During this time, mice were required to walk in a forward direction on the rotating dowels for as long as possible. When the mice were no longer able to walk on the rotating dowels, they fell onto the landing platform below. This triggered the end of the trial for an animal and measurements of time to fall (TTF) were collected. Passive rotations where mice clung to and consequently rotated with the dowel were also used to define the end of the trial. Mice were then returned to their cages with access to food and water for 10 min. This procedure was repeated for a total of six trials, with the first three trials used for training and subsequent trials used for data analysis.

### Forelimb grip strength

The forelimb grip strength of mice was measured using a grip strength meter (Columbus instrument, USA). Mice were held by the base of the tail close to the horizontal bar to allow them to reach and grab onto the bar with their forelimbs. Mice were then positioned so that their body was horizontal and in line with the bar. They were then pulled horizontally away from the bar by the tail until their grip was released. The tension was measured and defined as grip strength. Mice were given 1 min inter-trial intervals (ITI) during which they were returned to their cages with access to food and water. This procedure was repeated for a total of nine trials for each mouse (with the mean value of nine trials used for analysis).

### Frailty index assessment

The Frailty Index (FI) was assessed as described previously^25^. In brief, 31 health-related deficits were assessed for each mouse. A mouse was weighed, and body surface temperature was measured three times with an infrared thermometer (Fisher scientific). Bodyweight and temperature were scored based on their deviation from the mean weight and temperature of young mice^25^. Twenty-nine other items across the integument, physical/musculoskeletal, ocular/nasal, digestive/urogenital, and respiratory systems were scored as 0, 0.5, and 1 based on the severity of the deficit. The total score across the items was divided by the number of items measured to give a frailty index score between 0 and 1.

### MicroCT scan

Both femurs and tibia of the mice were analyzed in a Micro Computed Tomography (micro-CT) Facility (Tissue Imaging (BBMTI) Core in Center for Musculoskeletal Research, University of Rochester). Micro-CT was performed with a state-of-the-art scanner (VivaCT 40, Scanco USA, Inc.) for live small animals and specimens, without contrast agents. The scanner was fitted with an adjustable X Ray Source Energy (30 - 70 kVp) and scan (using Scancocone beam geometry) specimens in a field of view (FOV) of up to 39 mm and a scan length of 145 mm at a nominal resolution of 10 microns. Scan acquisition, reconstruction, analysis, and measurements are performed using a specialized suite of 64 bit software applications running on an open VMS platform.

The relevant 3D-images were imported after processing into the scan software and the parameters such as bone mass density (BMD), bone volume/tissue volume (BV/TV), bone surface/bone volume (BS/BV), bone surface/total volume (BS/TV), trabecular number (Tb.N), Trabecular separation (Tb-Sp) and connectivity density (Conn.Dn) were measured.

### Methylation clock

Genomic DNA from 24 months old CreER and nmrHAS2 mouse livers was purified using a DNeasy Blood & Tissue Kit (Qiagen). One hundred nanograms of purified genomic DNA were used for the methylation measurement. All DNAm data used was generated using the custom Illumina chip “HorvathMammalMethylChip40” so-called the mammalian methylation array. The particular subset of species for each probe is provided in the chip manifest file can be found at Gene Expression Omnibus (GEO) at NCBI as platform GPL28271. The SeSaMe normalization method was used to define beta values for each probe^81^. The methylation age was normalized to chronological age to calculate the age acceleration value.

To investigate the CpG sites that drive the epigenetic age difference between experimental and control groups, we tested the methylation levels of CpG sites reported to change during mouse aging in a previous study^24^. Paired Student t-test was used to calculate the statistical significance.

### 7,12-Dimethylbenz(a)anthracene/12-O-tetradecanoylphorbol-13-acetate (DMBA/TPA) treatment

Thirteen young CreER and eleven nmrHAS2 mice were topically treated with DMBA and TPA. A single dose of DMBA (7.8 mM dissolved in acetone) was topically treated to mice on the dorsal trunk. Three days after, 0.4 mM of TPA was treated 3 times per week. The formation of papilloma was quantified 20 weeks after TPA treatment.

### Measurement of cytokines and chemokines in plasma

Thirty-six cytokines and chemokines were measured in young and old mouse plasma by luminex multiplex technique using a Cytokine & Chemokine 36-Plex Mouse ProcartaPlex™ Panel 1A kit (Thermo Fisher Scientific). The luminex multiplex assay was performed using undiluted plasma samples following the manufactures instruction.

### Tissue section, immunofluorescence, and RNAscope *in situ* hybridization

Paraffin-embedded specimens were sectioned at 10 *μ*m thickness. Tissue sections were deparaffinized with xylene and dehydrated in a descending alcohol series of 100, 95, 80, 70, and 50%. These initial processing steps were the same for all the staining procedures described below and all staining procedures were performed on the same samples, samples were rehydrated in PBS for 30min before performing immunofluorescence. For antibody-based assays, the sections were incubated twice in antigen retrieval buffer (0.1 M sodium citrate, 0.1 M citric acid, pH 6.0) for 15 min at 90-100°C before blocking. All slides were blocked in TBS-T which contains 5% FBS and 1% BSA for 2 h at room temperature. Subsequently, sections were incubated overnight at 4°C with the following primary antibodies: Anti-MUC2 (1:1500, GeneTex), Anti-Lyzozyme (1:500, Abcam). Following the primary antibody incubation, the slides were washed three times in PBS-T and incubated with goat anti rabbit IgG (H+L) secondary antibody conjugated with Alexa Fluor 568(1:1000, Invitrogen) for 1h at room temperature. After a 5 times wash, slides were stained with DAPI (BioLegend) for 1min at room temperature, mounted with mounting media (Vector Laboratories) and observed under the confocal microscope at x40 magnification.

For HABP staining, after deparaffinization, slides were rehydrated in PBS for 30min at room temperature then blocked in TBS-T which contains 5% FBS and 1% BSA for 2h at room temperature. Then all slides were incubated with biotinylated hyaluronan binding protein (1:100 for small intestine and 1:200 for other tissues, Amsbio) overnight at 4°C. Following the HABP incubation, slides were washed and incubated with Streptavidin conjugated with Alexa Fluor 647(1:500, Thermo Fisher Scientific) for 1h at room temperature. Then the slides will be washed, counterstained with DAPI, mounted, and observed under the confocal microscope at x40 magnification. At least three random fields of each sample were captured for quantification of fluorescence signals. The average intensities of HA signals were quantified using ImageJ. Experiment was repeated from at least three animals of each group to confirm the reproducibility. RNAscope assay was performed using RNAscope Multiplex Fluorescent Detection Kit v2 according to the manufacturer’s protocol. All pictures were required under the confocal microscope at x40 magnification.

### Preparation of RNA for RT-qPCR and RNA sequencing

All frozen tissues were pulverized using the cell crusher. For preparing RNA from tissues, pulverized frozen tissues in the range of 10-15 mg were removed from samples kept at *−*80°C and extracted using Trizol reagent according to the supplier’s instructions. After recovery of total RNA from the Trizol reagent by isopropanol precipitation, RNA was digested with DNaseI for 30min at room temperature and further purified by RNeasy plus mini kit according to the instruction. For purifying RNA from cells, the RNeasy plus mini kit was used according to the user manual. The yield and quality were checked using Nano Drop.

### Reverse transcription quantitative PCR (RT-qPCR)

For RT-qPCR, ∼300ng of purified RNA was revers transcribed into cDNA in the 20 *μ*L using iScript cDNA synthesis kit (Bio-Rad). 2 *μ*L of this reaction was used for subsequent qPCR reactions, which were performed using SYBR green system (Bio-Rad). The primer sequences were listed in supplementary table 1. The actin beta gene was used as internal normalization control.

### RNA sequencing

The RNA samples were processed with the Illumina TruSeq stranded total RNA RiboZero Gold kit and then subjected to Illumina HiSeq 4000 single-end 150bp sequencing at New York University Genome Technology Center. Over 50 million reads per sample were obtained. The RNA-seq experiment was performed in three biological replicates for all tissues.

The RNA-seq reads were first processed using Trim_Galore (version 0.6.6), which trimmed both adapter sequences and low-quality base calls (Phred quality score < 20). The clean RNA-seq reads were used to quantify the gene expression with Salmon (version 1.4.0)^82^. Specific parameters (--useVBOpt --seqBias --gcBias) were set for sequence-specific bias correction and fragment GC bias correction. Gencode^83^ (version M25) was used for the genome-wide annotation of the gene in the mouse. The reads counts for genes were used as the input for differential expression analysis by DESeq2^84^. Low-expression genes with reads counts less than 10 were excluded. The cutoff for p-value and fold change was shown in the figure or figure legend.

For the hierarchical clustering analysis, we reanalyzed published gene expression data of mice treated with rapamycin, calorie restriction, growth hormone knockout, etc. To minimize the potential batch effect, we directly compared the gene expression data of mice with intervention treatments to its corresponding control. Fold changes (intervention / control) of gene expression level were calculated for each gene. Log2 scaled fold changes of gene expression were used to perform the analysis.

Gene set enrichment analysis (GSEA)^85^ was performed with “Preranked” model (version 4.1.0). All genes were preranked by the values of –log10 (adjusted p value)*(fold change)/abs(fold change). Adjusted p values and fold change were obtained from DEseq2. Normalization mode was set to “meandiv”. Only those gene set with a size more than 15 genes were kept for the further analysis.

GO analysis were performed by R package clusterProfiler (Release version 3.14)^86^. GO comprises of three orthogonal ontologies, i.e. molecular function (MF), biological process (BP), and cellular component (CC). All the p-values are adjusted following Benjamini –Hochberg (BH) correction.

Association of gene expression log-fold changes induced by nmrHas2 expression in mouse livers with previously established transcriptomic signatures of aging and lifespan-extending interventions was examined as described in the previous study^29^ separately for males and females. Utilized signatures of aging included tissue-specific brain, liver and kidney signatures as well as multi-tissue signatures of mouse, rat and human. Signatures of lifespan-extending interventions included genes differentially expressed in mouse tissues in response to individual interventions, including caloric restriction (CR), rapamycin (Rapamycin), and mutations associated with growth hormone deficiency (GH deficiency), along with common patterns of lifespan-extending interventions (Common) and expression changes associated with the mouse maximum (Max lifespan) and median lifespan (Median lifespan).

Pairwise Spearman correlation between logFC induced by nmrHas2 expression and associated with signatures of aging and longevity was calculated based on the union of top 400 statistically significant genes (with the lowest p-value) for each pair of signatures.

For the identification of enriched functions affected by nmrHas2 expression in mouse livers we performed Fisher exact test and functional GSEA (<u>10.1073/pnas.0506580102</u>) on a pre-ranked list of genes based on log_10_(p-value) corrected by the sign of regulation, calculated as:

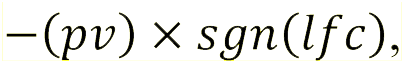

where *pv* and *lfc* are p-value and logFC of a certain gene, respectively, obtained from edgeR output, and *sgn* is the signum function (equal to 1, -1 and 0 if value is positive, negative or equal to 0, respectively). HALLMARK, KEGG and REACTOME ontologies from the Molecular Signature Database (MSigDB) were used as gene sets. Fisher exact test and GSEA were performed separately for each sex using *gprofile2* and *fgsea* R packages, respectively. A q-value cutoff of 0.1 was used to select statistically significant functions.

Similar analysis was performed for gene expression signatures of aging and lifespan-extending interventions. Pairwise Spearman correlation was calculated for individual signatures of nmrHas2 expression, aging and lifespan-extending interventions based on NES estimated with GSEA. A heatmap colored by NES was built for manually chosen statistically significant functions (adjusted p-value < 0.1). Complete list of functions enriched by genes perturbed by nmrHas2 expression in mouse livers is included in Extended Data Table 3.

### Primary fibroblast isolation and cell culture

Primary skin fibroblasts were isolated from under-arm skin from 5 months old CreER and nmrHAS2 mice. Skin tissues were shaved and cleaned with 70% ethanol then minced and incubated in DMEM/F-12 medium (ThermoFisher) with Liberase (Millipore Sigma) at 37*℃* on a stirrer for 40min. Tissues were then washed and plated with DMEM/F-12 medium containing 15% fetal bovine serum (GIBCO) and Antibiotic-Antimycotic (GIBCO). When cells are 80% confluent, isolated cells were frozen in liquid nitrogen within 2 passages. All subsequent fibroblast cultures were performed in EMEM (ATCC) supplemented with 10% fetal bovine serum (GIBCO), 100 units/mL penicillin, and 100 mg/mL streptomycin (GIBCO). Raw264.7 cell line was purchased from ATCC and maintained using DMEM (Gibco) supplemented with 10% fetal bovine serum (GIBCO), 100 units/mL penicillin, and 100 mg/mL streptomycin (GIBCO). The Raw cells used for all experiments are all under passage four. All primary cells were cultured at 37*℃* with 5% CO_2_ and 3% O_2_.

### Apoptosis assay

Skin fibroblast cells under population doubling 15 were used for H_2_O_2_ treatment. Cells with ∼80% confluency were treated with H_2_O_2_ at concentrations of 100 µM, 200 µM, and 400 µM for 24h. Cells were collected, and apoptotic cells were quantified using Annexin V FLUOS Staining Kit (Roche) following the manufacturer’s instructions. After staining, cells were analyzed with CytoFlex flow cytometer (Beckman). Cells which are double negative for Annexin-V and PI signals were defined as live cells.

### Bone marrow derived macrophage isolation, culture, and LPS challenge

Bone marrow derived macrophages were isolated from 5 months old CreER and nmrHAS2 mice. Mice were sacrificed via cervical dislocation and hind legs were dissected. Using aseptic technique, bone marrow was extracted from tibia and femur bones following removal of surrounding muscle. To do so, joints were cut using a scalpel and the exposed bone marrow was flushed out the ends of the bones using a 27-gauge needle and a 10 ml syringe filled with cold RPMI-1640 media. Clumps were gently disaggregated using a needle-less syringe and passed through a 70 *μ*m cell strainer. The cell suspension was centrifuged at 250 g for 5 min at room temperature to pellet cells. Bone marrow cells were subsequently cultured in RPMI-164 (GIBCO) supplemented with 10% fetal bovine serum (GIBCO), 100 units/mL penicillin, 100 mg/mL streptomycin (GIBCO), and 10ng/mL recombinant mouse M-CSF(R&D) at 37*℃* with 5% CO_2_ and 3% O_2_ on Day 0. Fresh media were changed every 48h, Double volume of media were used on Day 4 and M-CSF was supplemented on Day 6 to avoid removing the HA produced by cells. 10ng/mL LPS was used to treat macrophages on Day 7. Cells were harvest 24h after LPS treatment for RNA extraction.

### Gut permeability assay

Tracer FITC-labeled dextran (4kDa; Sigma-Aldrich) was used to assess in vivo intestinal permeability. Mice were deprived of food 8 h prior to and both food and water following an oral gavage using 200 ml of 80 mg/ml FITC-dextran. Blood was retro-orbitally collected after 4 h, and fluorescence intensity was measured on fluorescence plates using an excitation wavelength of 493nm and an emission wavelength of 518 nm. The untreated mouse plasma was used as blank.

### Isolation and culture of primary intestinal crypts

Around 20 cm small intestines from 7- and 18-months old mice were removed and flushed using 10 mL syringe with clear lumenal contents. Then the intestines were opened longitudinally, washed in 20 mL cold PBS and cut into ∼5 mm sections, and placed into the 50 mL canonical tube which contains 15 mL cold PBS. Intestinal crypts were mechanically released from the lamina propria by vigorous pipetting and washed using PBS for twenty times followed by incubating in 25 mL gentle cell dissociation reagent (STEMCELL Technology) for 15min at room temperature. After that, the dissociation reagent was neutralized by adding 10 mL of cold PBS containing 1% BSA. The supernatants containing crypts were filtered through a 70-µm cell strainer and centrifuged at 290 xg for 5 min at 4°C then washed one more time with cold PBS containing 1% BSA. Isolated crypts were counted and embedded in Matrigel (Corning) on the ice at ∼12.5 crypts per µl and mixed with the same volume of IntestiCult Organoid Growth medium (STEMCELL Technology). Matrigel beads occupying the center of the well were constructed using 40 µl of Matrigel to form a solid dome-like structure in an 8-well chamber slide and were subsequently overlaid with 400 µl IntestiCult Organoid Growth medium. For the HA treatment, 20 µg/mL HMW or LMW HA (R&D) were directly added to the Matrigel crypts mixture. Primary intestinal crypts were incubated in a fully humidified culture chamber with 5% CO_2_ at 37°C. The culture medium was changed every 48 h, and the organoid-forming efficiency was calculated on days 4.

### Microbiota analysis

Fresh fecal samples from 9- and 23-months old animals were obtained in the morning, immediately snap freeze in liquid nitrogen, and stored in -80*℃*. These samples were used for 16S rRNA gene analysis for microbiota profiling from the V1–V2 region of 16S rRNA genes. DNA extraction was performed using QIAamp DNA Stool Mini Kit (Qiagen) following the manufacture’s instruction. DNA was eluted in 50 µl DNAse free water. Twenty nanograms of DNA were used for the amplification of the 16S rRNA gene with primers 27F-DegS (5’-TCGTCGGCAGCGTCAGATGTGTATAAGAGACAGGTTYGATYMTGGCTCAG-3’) and 338R I (5’-GTCTCGTGGGCTCGGAGATGTGTATAAGAGACAGGCWGCCTCCCGTAGGAGT-3’) + 338R II (5’-GTCTCGTGGGCTCGGAGATGTGTATAAGAGACAGGCWGCCACCCGTAGGTGT-3’)^87^ for 25 cycles. Primers have Illumina sequencing index attached; Index forward (5’-TCGTCGGCAGCGTCAGATGTGTATAAGAGACAG-3’) and index reverse (5’-GTCTCGTGGGCTCGGAGATGTGTATAAGAGACAG-3’). The PCR was performed in a total volume of 50 µl containing 1× HF buffer (New England BioLabs), 1 µl dNTP Mix (New England BioLabs), 1 U of Phusion® Hot Start II High-Fidelity DNA polymerase (New England BioLabs), 500 nM of the 27F-DegS primer, 500 nM of an equimolar mix of two reverse primers, 338R I and II. The size of the PCR products (∼375 bp) was confirmed by gel electrophoresis using 5 µl of the amplification reaction mixture on a 1% (w/v) agarose gel. The PCR products were purified using QIAquick PCR Purification Kit (Qiagen). Then, the dual indices and Illumina sequencing adapters were attached using the Nextera XT index kit (Illumina) following the manufacture’s instruction. Purified amplicon pools were 250 bp paired end sequenced using Illumina Miseq.

The Illumina Miseq data analysis was carried out with a workflow employing the Quantitative Insights Into Microbial Ecology (QIIME2) pipeline^88^. The reads were processed the as following: reads were filtered for not matching barcodes; OTU picking and chimera removal was done via matching the sequences to the Silva 111 database, with only one mismatch allowed, and a biom and with clustalw a multiple alignment and phylogenetic tree file was generated. Further outputs were generated via QIIME, such as filtered reads per sample, PD whole tree diversity measurements and the level 1–6 taxonomic distributions with relative abundances. 37,000 reads cutoff was used for all the samples.

### Statistical and demographic analysis

Data are shown as means with SEM (unless stated otherwise). N indicates the number of animals per test group; age and sex are also noted. Student’s t test (unpaired, two-tailed, equal variance) was used for all pairwise comparisons which satisfied with normal distribution. Mann-Whitney u test was used for data which is not satisfied with normal distribution. All relevant p values are shown in the figures; if not shown, * indicates p<0.05, ** indicates p<0.01, *** indicates p<0.001, **** indicates p<0.0001 and ns means no significance. Demographic data were processed with GraphPad Prism software to compute mean and median lifespans, SEM, percent increase of the median, and p values (log-rank test) for each cohort.

---

**Extended data Figure 1.**
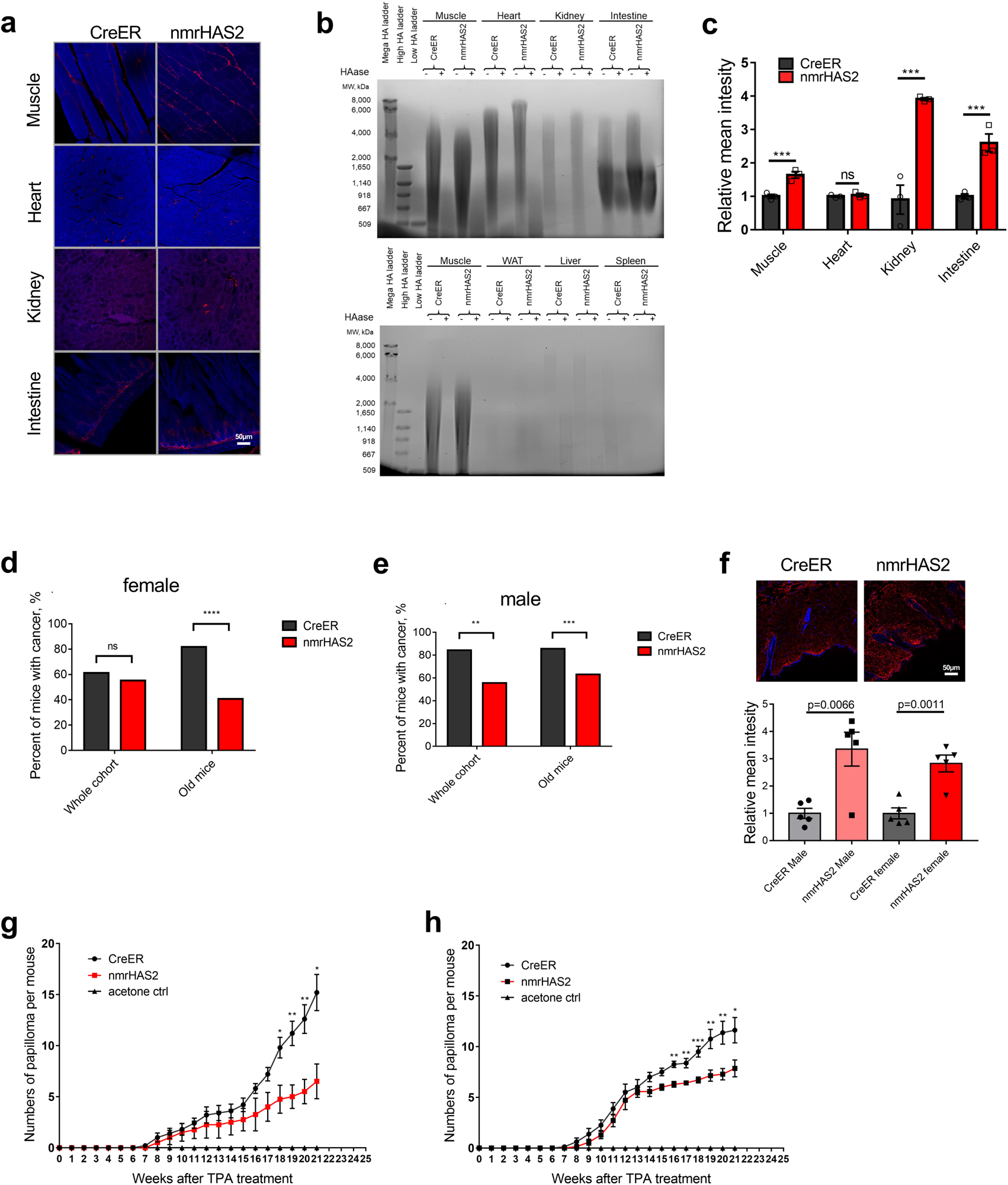
nmrHAS2 mice exhibit resistance to spontaneous and induced cancer. **a.** Representative pictures of HABP staining in organs of male nmrHAS2 and CreER mice. **b.** Pulse field gel shows that male nmrHAS2 mice have higher molecular weight and more abundant hyaluronic acid. HA was extracted from 200 mg of pooled tissue from two individuals. HAase treated samples were run in parallel to confirm the specificity of HA staining. **c.** Levels of relative HABP fluorescence intensity. N=3. *** p < 0.001, unpaired Student t-test. **d.** Old female nmrHAS2 mice have much lower spontaneous cancer incidence (N=47-49). **** p < 0.0001, p-values were calculated using Chi-square test. **e.** Old male nmrHAS2 mice have much lower spontaneous cancer incidence (N=27-32). ** indicates p<0.01, *** indicates p<0.001, p-values were calculated using Chi-square test. **f.** HABP staining shows that nmrHAS2 mice skin has a higher hyaluronan level. N=5, unpaired Student t-test. **g.** Quantification of papilloma formation in DMBA/TPA treated female mice (N=3-5). * indicates p<0.05, ** indicates p<0.01, p-values were calculated using two-tailed Student’s t-test. **h.** Quantification of papilloma formation in DMBA/TPA treated male mice (N=5-8). * indicates p<0.05, ** indicates p<0.01, *** indicates p<0.001, p-values were calculated using two-tailed Student’s t-test.

**Extended data Figure 2.**
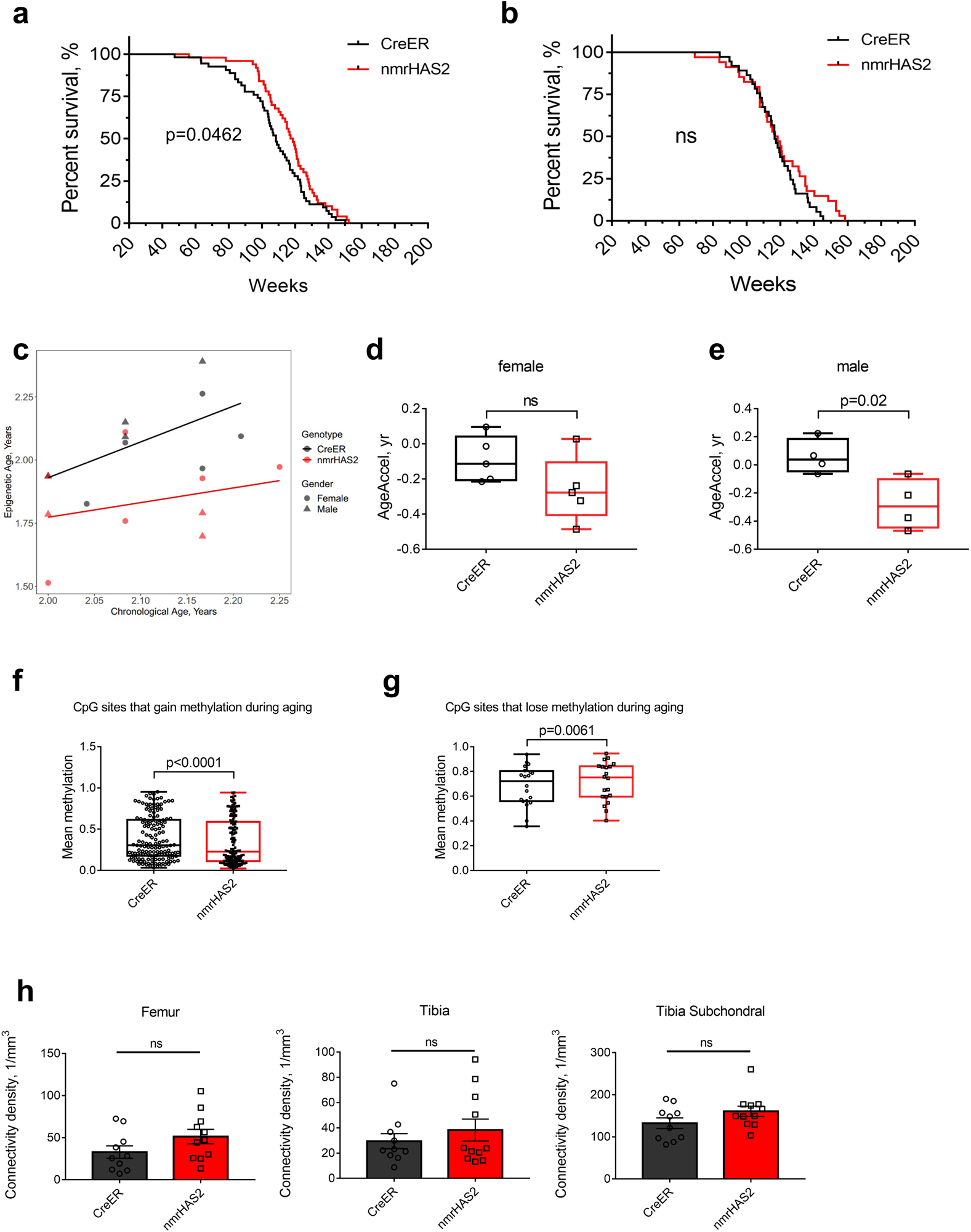
nmrHAS2 mice showed extended lifespan and healthspan. **a.** Female nmrHAS2 mice have extended median lifespan (N=50-53). p-value for median lifespan was calculated using log-rank test. **b.** Male nmrHAS2 mice have extended maximum lifespan (N=31-37). p-value for median lifespan was calculated using log-rank test. **c.** Old nmrHAS2 mice display younger epigenetic age. **d.** Old nmrHAS2 female mice display younger biological age. p-values were calculated using unpaired Student’s t-test. **e.** Old nmrHAS2 male mice display younger biological age. p-values were calculated using unpaired Student’s t-test.

**Extended data Figure 3.**
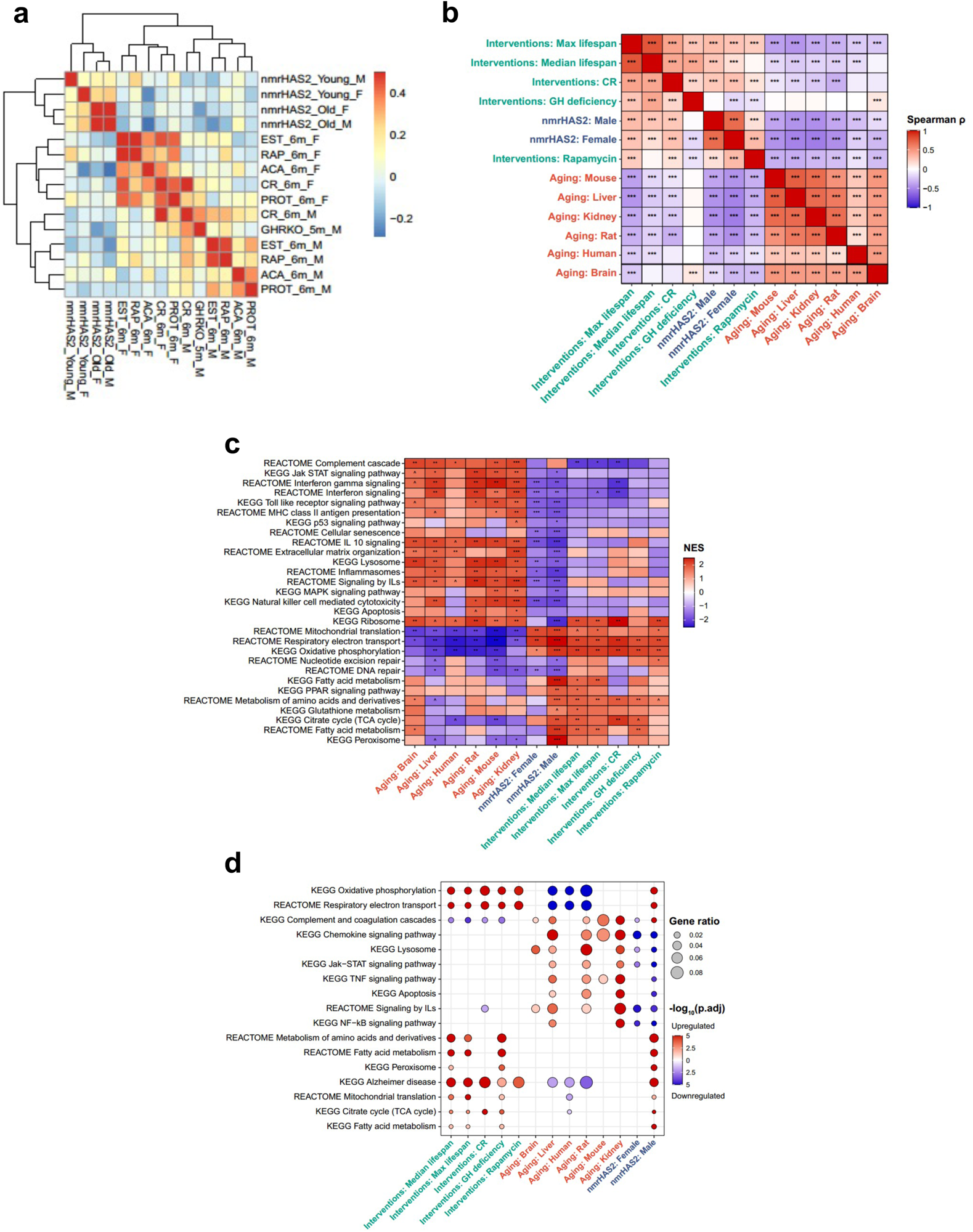
nmrHAS2 mice showed a distinct transcriptomic signature. **a.** nmrHAS2 mice display an expression signature distinct form mice subjected to other pro-longevity interventions. A heatmap of correlation analysis performed on liver whole transcriptomes. **b.** Association between nmrHAS2 effect and signatures of lifespan-extending interventions and mammalian aging based on functional enrichment (GSEA) scores. Only functions enriched by at least one signature (adjusted p-value < 0.1) were used for the calculation. **c.** Functional enrichment (GSEA) of gene expression signatures associated with nmrHAS2, mammalian aging and established lifespan-extending interventions. Only functions significantly enriched by at least one signature (adjusted p-value < 0.1) are presented. **d.** Functional enrichment (Fisher exact test) of genes significantly associated with the effect of nmrHAS2, mammalian aging and established lifespan-extending interventions. Only functions enriched by at least one aggregated signature (adjusted p-value < 0.1) are shown. Proportion of pathway-associated genes is reflected by bubble size. ^ p.adjusted < 0.1; * p.adjusted < 0.05; ** p.adjusted < 0.01; *** p.adjusted < 0.001.

**Extended data Figure 4.**
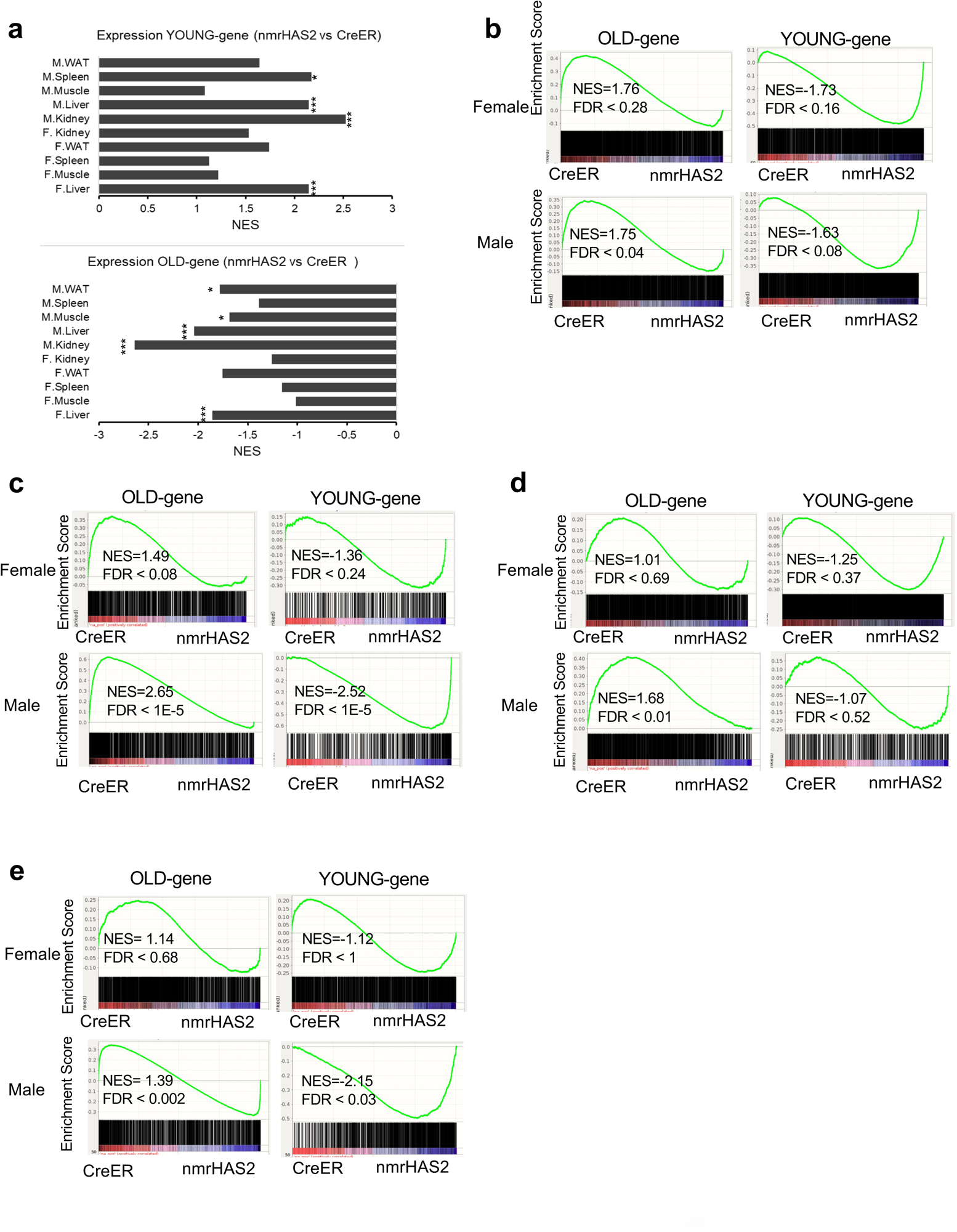
nmrHAS2 mice showed a younger transcriptomic state. **a.** GSEA plots show that YOUNG gene set is upregulated, and OLD gene set is downregulated in all sequenced old nmrHAS2 mice for both sexes. * FDR<0.05, *** FDR<0.001. **b.** GSEA plots showing that OLD gene set is downregulated in WAT of old male nmrHAS2 mice. **c.** GSEA plots showing that that YOUNG gene set is upregulated, and OLD gene set is downregulated in the kidney of old male nmrHAS2 mice. **d.** GSEA plots showing that OLD gene set is downregulated in the muscle of old male nmrHAS2 mice. **e.** GSEA plots showing that that YOUNG gene set is upregulated, and OLD gene set is downregulated in the spleen of old male nmrHAS2 mice.

**Extended Data Figure 5.**
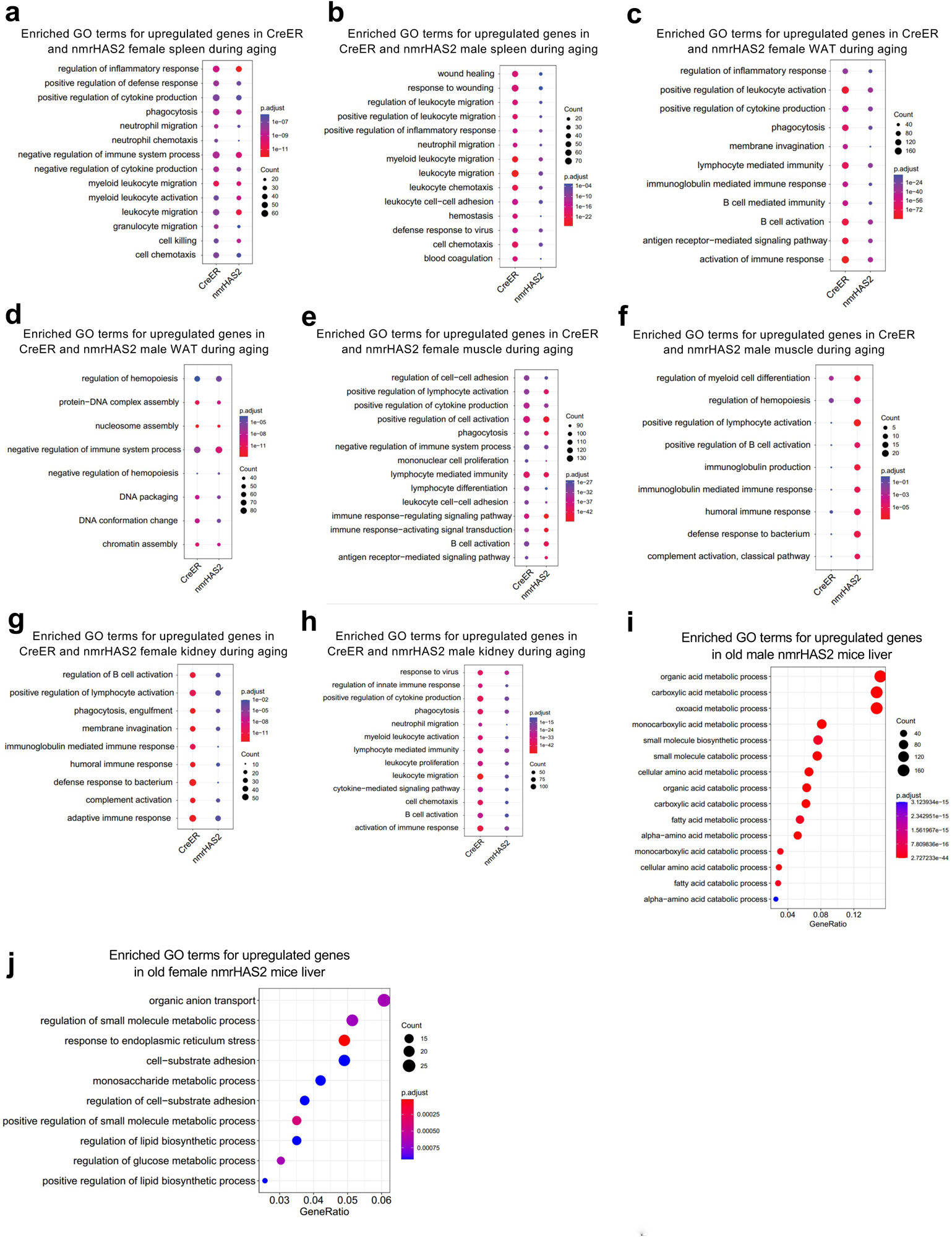
RNAseq shows nmrHAS2 mice have reduced inflammation during aging. **a.** Enriched GO terms for upregulated genes in the spleen of CreER and nmrHAS2 female mice during aging. **b.** Enriched GO terms for upregulated genes in the spleen of CreER and nmrHAS2 male mice during aging. **c.** Enriched GO terms for upregulated genes in the WAT of CreER and nmrHAS2 female mice during aging. **d.** Enriched GO terms for upregulated genes in the WAT of CreER and nmrHAS2 male mice during aging. **e.** Enriched GO terms for upregulated genes in the muscle of CreER and nmrHAS2 female mice during aging. **f.** Enriched GO terms for upregulated genes in the muscle of CreER and nmrHAS2 male mice during aging. **g.** Enriched GO terms for upregulated genes in the kidney of CreER and nmrHAS2 females during aging. **h.** Enriched GO terms for upregulated genes in CreER and nmrHAS2 male kidney during aging. **i.** Enriched GO terms for upregulated genes in old male nmrHAS2 mice liver **j.** Enriched GO terms for upregulated genes in old female nmrHAS2 mice liver

**Extended Data Figure 6.**
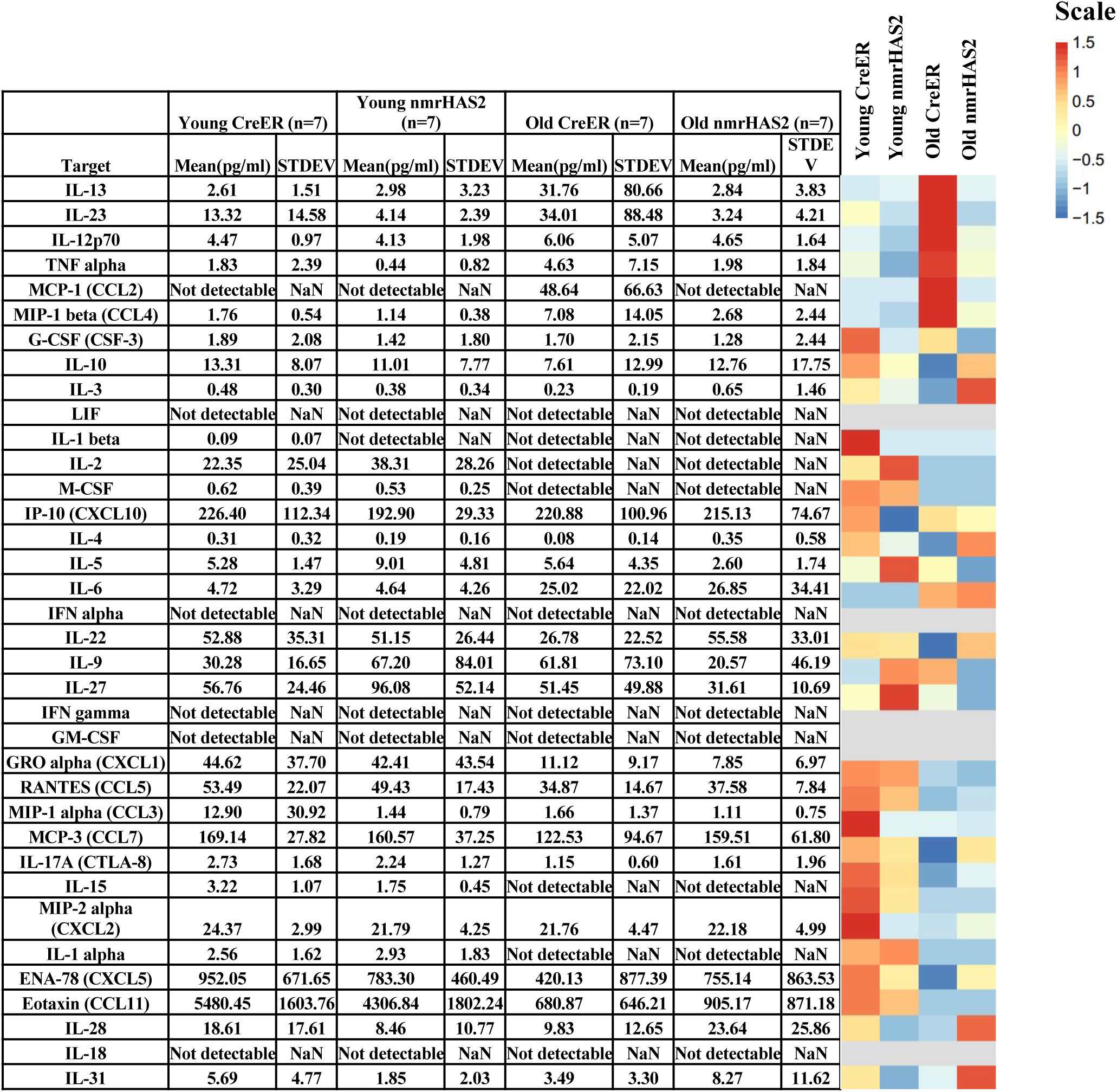
Mean plasma concentrations of 36 inflammatory cytokines in nmrHAS2 and CreER male mice. Mean plasma concentrations of 36 inflammatory cytokines and chemokines of young (5-months) and old (24-months) male mice. The heatmap is presented alongside the value chart. In the heatmap, the level of each target was scale automatically using R.

**Extended Data Figure 7.**
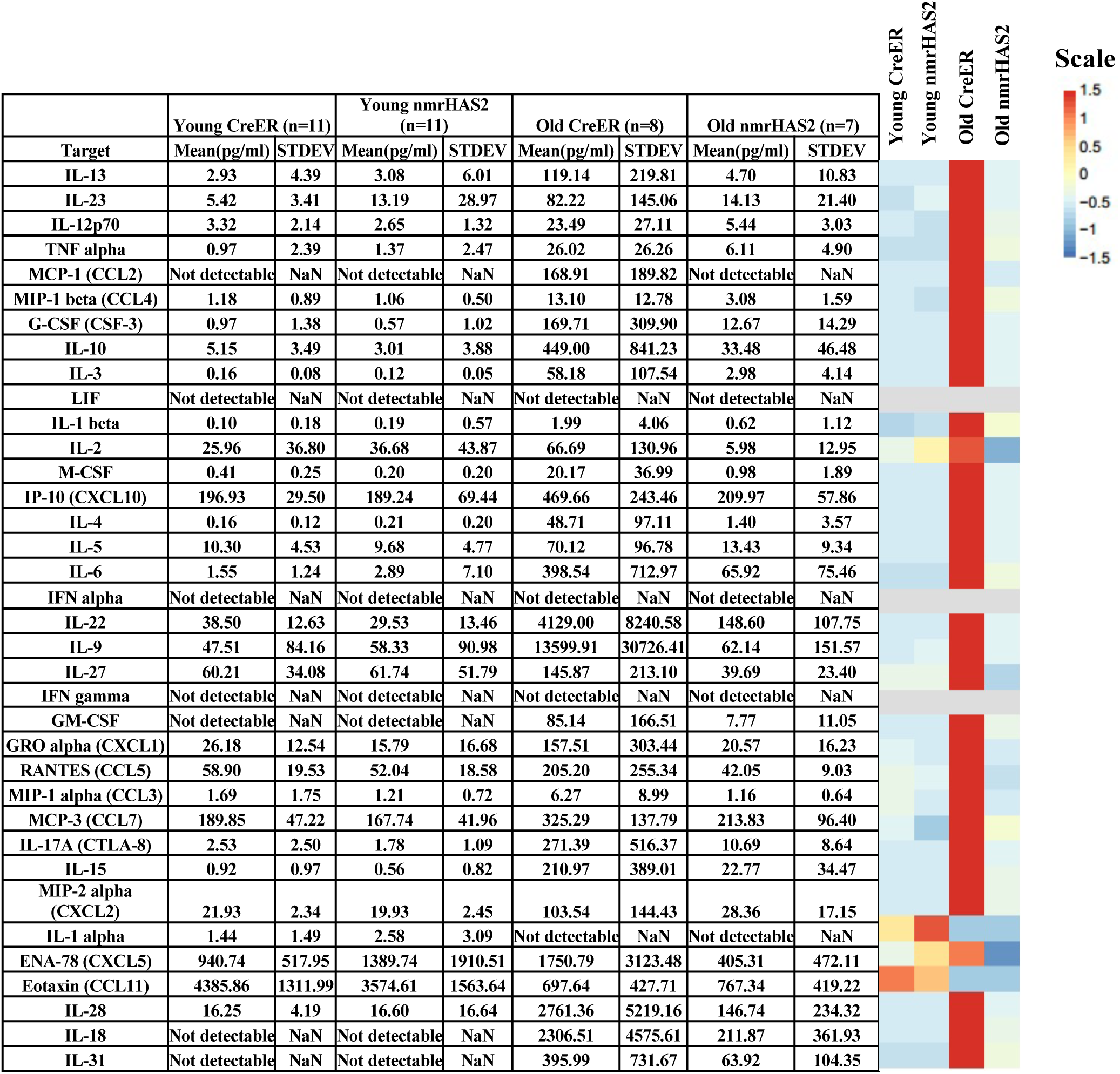
Mean plasma concentrations of 36 inflammatory cytokines in nmrHAS2 and CreER female mice. The mean plasma concentrations of 36 inflammatory cytokines and chemokines in young (5-months) and old (24-months) female mice. The heatmap is presented alongside the value chart. In the heatmap, the level of each target was scaled automatically using R.

**Extended data Fig. 8.**
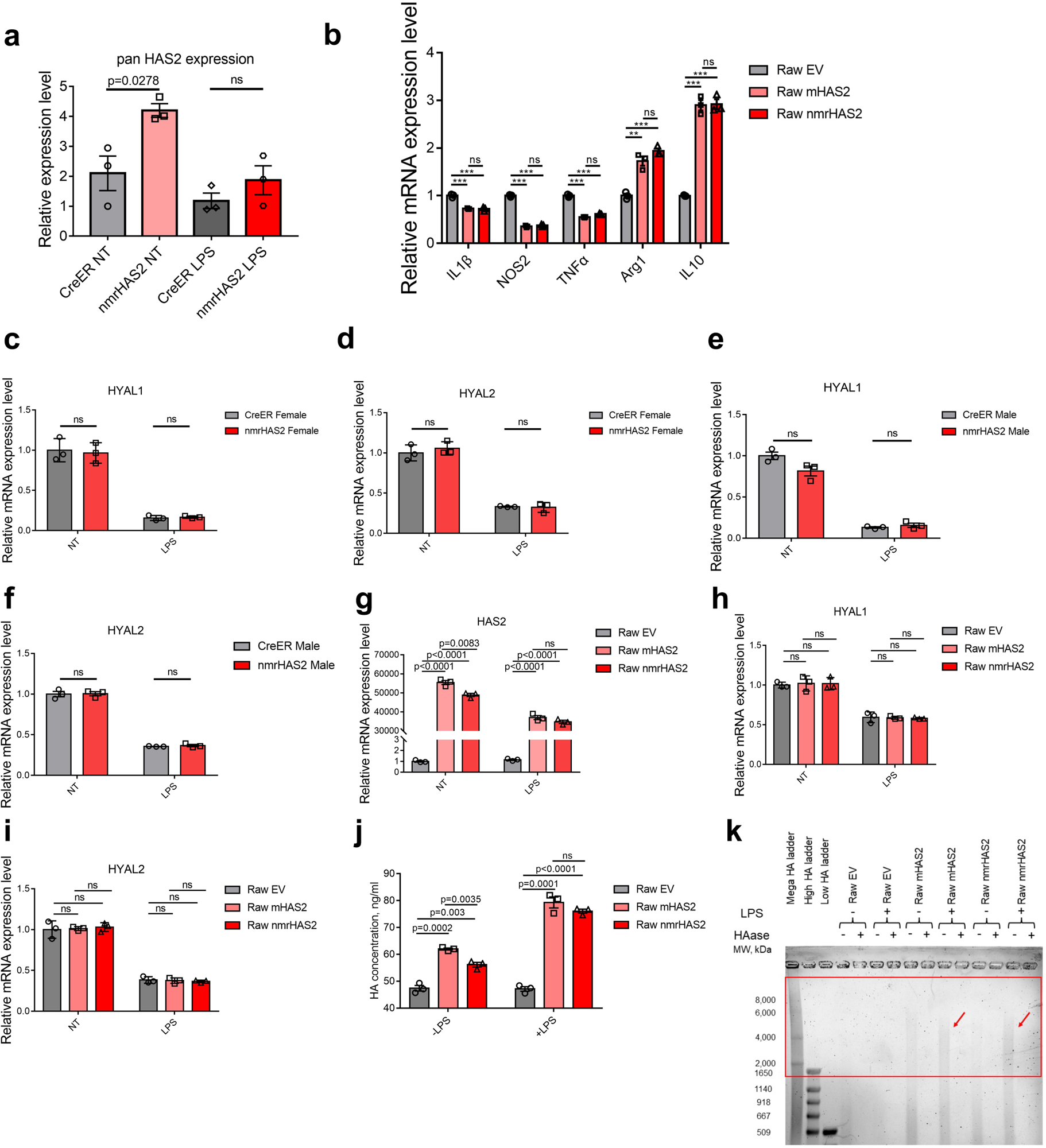
nmrHAS2 reduces pro-inflammatory response *in vitro*. **a.** BMDM from nmrHAS2 mice have significantly upregulated HAS2 levels. BMDM were isolated from 5-months old female mice (N=3). p-values were calculated using two-tailed Student’s t-test. **b.** Raw264.7 cells overexpressing mHAS2 or nmrHAS2 show lower levels of pro-inflammatory cytokines and higher levels of anti-inflammatory cytokines. Data was normalized to Raw EV. p-values were calculated using unpaired Student’s t-test, N=3. **c.** HYAL1 levels decrease after LPS treatment in BMDM from female mice. Normalization to CreER NT. p-values were calculated using two-tailed Student’s t-test; N=3. **d.** HYAL2 levels decrease after LPS treatment in BMDM from female mice. Normalization to CreER NT. p-values were calculated using two-tailed Student’s t-test; N=3. **e.** HYAL1 levels decrease after LPS treatment in BMDM from male mice. Normalization to CreER NT. p-values were calculated using two-tailed Student’s t-test, N=3. **f.** HYAL2 levels decrease after LPS treatment in BMDM from male mice. Normalization to CreER NT. p-values were calculated using two-tailed Student’s t-test; N=3. **g.** HAS2 levels decrease in LPS treated HAS2 expressing Raw264.7 cells. Normalization to Raw EV NT. p-values were calculated using unpaired Student’s t-test; N=3. **h.** HYAL1 levels decrease after LPS treatment in HAS2 expressing Raw264.7 cells. Normalization to Raw EV NT. p-values were calculated using unpaired Student’s t-test; N=3. **i.** HYAL2 levels decrease after LPS treatment in HAS2 expressing Raw264.7 cells. Normalization to Raw EV NT. p-values were calculated using unpaired Student’s t-test; N=3. **j.** Raw264.7 cells overexpressing HAS2 produce more HA. HA ELISA was used to quantify the HA level in the media. p-values were calculated using two-tailed Student’s t-test; N=3. **k.** Raw264.7 cells overexpressing HAS2 produce more HMW-HA in the media after LPS treatment. Red square indicates the HMW-HA.

**Extended data Fig. 9.**
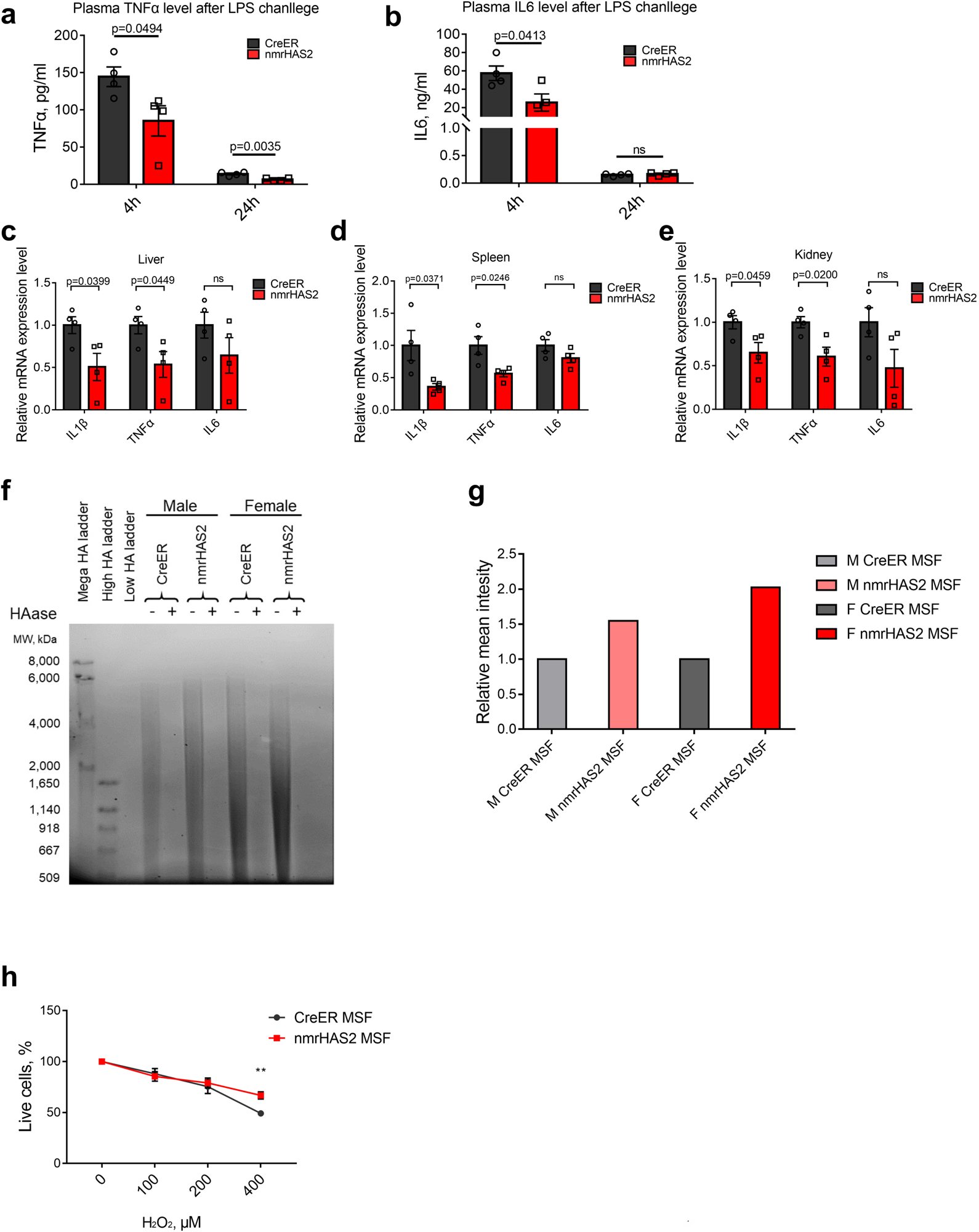
nmrHAS2 reduces pro-inflammatory response *in vivo* and protects cells from oxidative stress. **a.** nmrHAS2 mice produce significantly lower plasma TNFα levels 4 h and 24 h after LPS challenge in 5-months old female mice (N=4). p-values were calculated using two-tailed Student’s t-test. **b.** nmrHAS2 mice produced significantly lower plasma IL6 levels 4 h after LPS challenge in 5-months old female mice (N=4). p-values were calculated using two-tailed Student’s t-test. **c.** nmrHAS2 mice showed lower IL1β and TNFα levels in liver 24 h post LPS challenge in 5-months old female mice (N=4). p-values were calculated using two-tailed Student’s t-test. **d.** nmrHAS2 mice show lower IL1β and TNFα levels in the spleen 24 h post LPS challenge in 5-months old female mice (N=4). p-values were calculated using two-tailed Student’s t-test. **e.** nmrHAS2 mice show lower IL1β and TNFα levels in kidney 24 h post LPS challenge in 5-months old female mice (N=4). p-values were calculated using two-tailed Student’s t-test. **f.** Pulse field gel shows nmrHAS2 skin fibroblasts produce more hyaluronic acid. compared to CreER fibrobalsts. HAase treated samples were run in parallel to confirm the specificity of HA staining. Media from three different cell lines was pooled for HA extraction. **g.** Levels of relative on gel HA intensity. The intensity of HA was quantified using ImageJ. Intensity of nmrHAS2 group was normalized to the CreER group. **h.** Skin fibroblasts isolated from nmrHAS2 mice are more resistant to H_2_O_2_ treatment. Fibroblasts were isolated from 5-months old female mice (N=4). ** indicates p<0.01, p-values were calculated using two-tailed Student’s t-test.

**Extended data Fig. 10.**
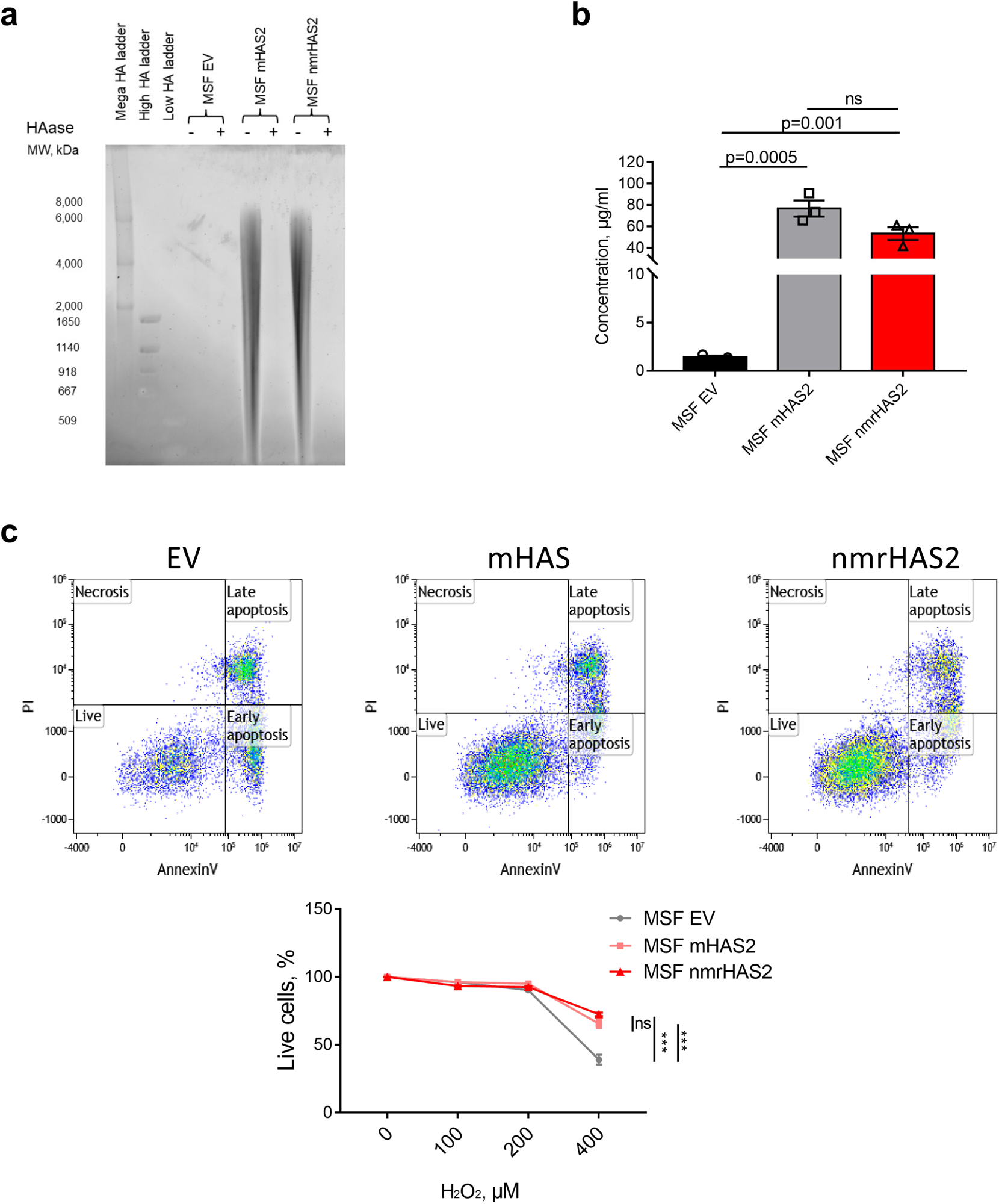
Overexpression of mouse or nmrHAS2 protects cells from oxidative stress. a. Pulse field gel shows that mouse skin fibroblasts (MSF) overexpressing mouse HAS2 (mHAS2) or nmrHAS2 produce more hyaluronic acid compared to fibroblasts transfected with empty vector (EV). HAase-treated samples were run in parallel to confirm the specificity of HA staining. Media from three different cell lines was pooled for HA extraction. b. HA ELISA shows that mouse skin fibroblasts (MSF) overexpressing mHAS2 or nmrHAS2 produce more hyaluronic acid compared to fibroblasts transfected with empty vector (EV). p-values were calculated using two-tailed Student’s t-test, N=3. c. Mouse skin fibroblasts overexpressing mHAS2 or nmrHAS2 are more resistant to H_2_O_2_ treatment. *** indicates p<0.001 p-values were calculated using two-tailed Student’s t-test, N=3.

**Extended data Figure 11.**
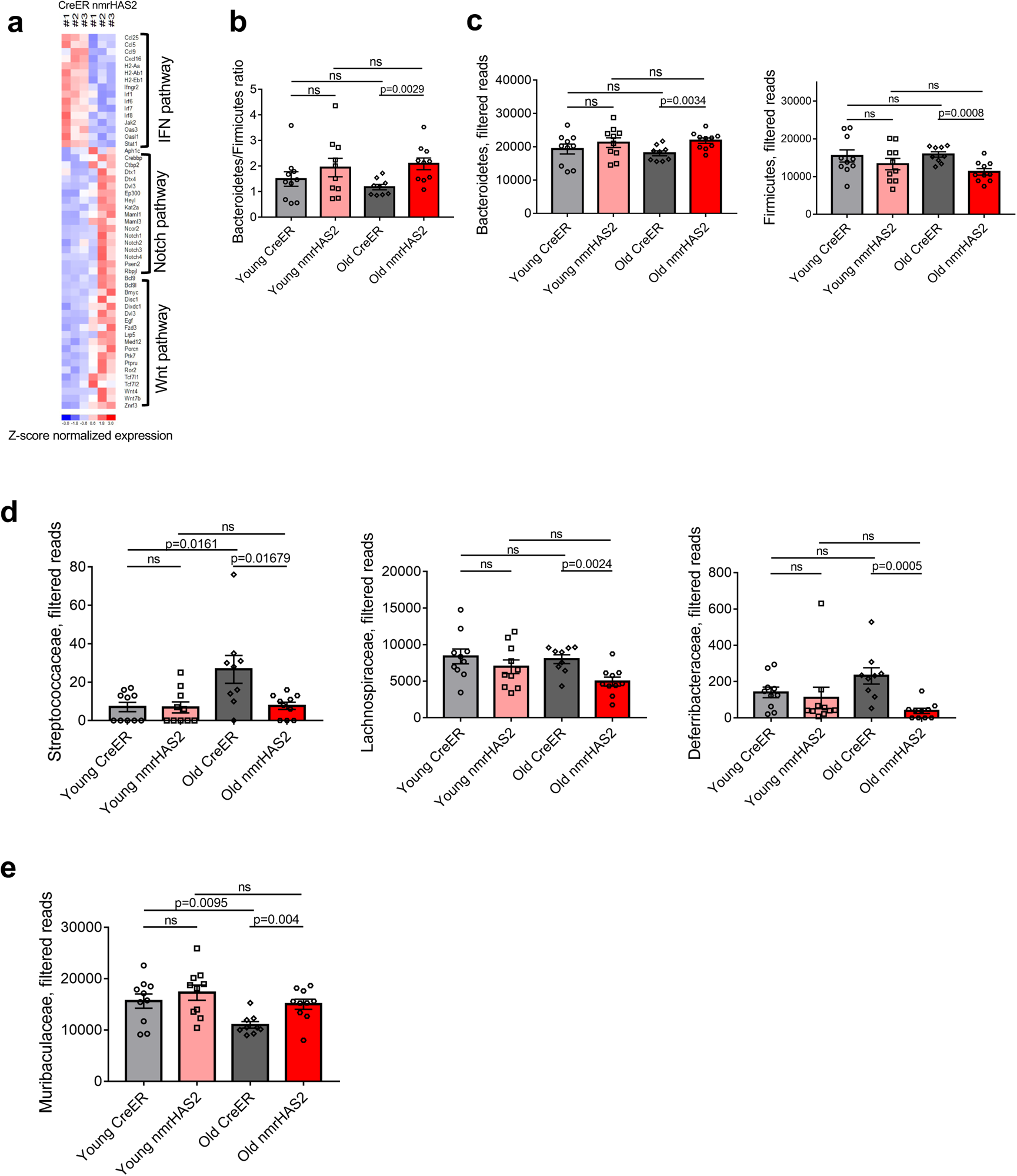
Old nmrHAS2 mice differ from age-matched CreER controls in their gut microbiome composition. **a.** Heatmap of genes involve in IFN, WNT, and Notch pathways. **b.** 6s rRNA sequencing shows old nmrHAS2 mice have a higher B/F ratio. Pooled females and males (n=9-10). p-values were calculated using two-tailed Student’s t-test. **c.** 16s rRNA sequencing shows that at the phylum level old nmrHAS2 mice have more abundant *Bacteroidetes* and less abundant *Firmicutes* compared to age-matched controls. 7- and 24-month-old mice were used. Pooled females and males (N=9-10). p-values were calculated using two-tailed Student’s t-test. **d.** 16s rRNA sequencing shows that at family level old nmrHAS2 mice have less abundant pro-inflammatory *Streptococcaceae*, *Lachnospiraceae,* and *Deferribacteraceae* compared to age-matched controls. 7- and 24-month-old mice were used. Pooled females and males (N=9-10). p-values were calculated using two-tailed Student’s t-test. **e.** 16s rRNA sequencing shows that at family level old nmrHAS2 mice have more *Muribaculaceae* compared to the age-matched controls. Pooled females and males (N=9-10). p-values were calculated using two-tailed Student’s t-test.

