## Extended Data Table 1 for "Increased hyaluronan by naked mole-rat HAS2 extends lifespan in mice"

Extended Data Table 1. nmrHAS2 mRNA level in different organs

| **Tissue** | **Gender** | **Log2(nmrHAS2/CreER)** | **p value** |
| --- | --- | --- | --- |
| Liver | Female | 7.7 | 2E-09 |
| Muscle | Female | 9.5 | 2E-136 |
| Spleen | Female | 3.9 | 3E-38 |
| WAT | Female | 1.8 | 3E-04 |
| Kidney | Female | 10.6 | 1.77E-15 |
| Intestine | Female | 5.7 | 9E-54 |
| Liver | Male | Not detectable | N/A |
| Muscle | Male | 8.7 | 1E-17 |
| Spleen | Male | 3.0 | 4E-05 |
| WAT | Male | 3.2 | 3E-08 |
| Kidney | Male | 10.4 | 8.02E-08 |
| Intestine | Male | 5.1 | 2E-22 |
