## Supplementary table 1 for "Increased hyaluronan by naked mole-rat HAS2 extends lifespan in mice"

**Supplementary Table 1. Primer sequences used for quantitative RT-PCR**

| Gene | Forward primer | Reverse primer |
| --- | --- | --- |
| *actb* | 5'-GGCTGTATTCCCCTCCATCG-3' | 5'-CCAGTTGGTAACAATGCCATGT-3' |
| *HAS2* | 5'-GCCTCATCTGTGGAGATGGT-3' | 5'-GAGTTTCTGTACATTCCCAGAGG-3' |
| *Il1b* | 5'-CCGTGGACCTTCCAGGATGA-3' | 5'-GGGAACGTCACACACCAGCA-3' |
| *Il6* | 5'-AGTTGCCTTCTTGGGACTGA-3' | 5'-TCCACGATTTCCCAGAGAAC-3' |
| *tnfa* | 5'-CATCTTCTCAAAATTCGAGTGACAA-3' | 5'-TGGGAGTAGACAAGGTACAACCC-3' |
| *arg1* | 5'-GCTCAGGTGAATCGGCCTTTT-3' | 5'-TGGCTTGCGAGACGTAGAC-3' |
| *Il12b* | 5'-AGACCCTGCCCATTGAACTG-3' | 5'-GAAGCTGGTGCTGTAGTTCTCATATT-3' |
| *nos2* | 5'-TTCACCCAGTTGTGCATCGACCTA-3' | 5'-TCCATGGTCACCTCCAACACAAGA-3' |
| *Il10* | 5'-GAGAGCTGCAGGGCCCTTTGC-3' | 5'-CTCCCTGGTTTCTCTTCCCAAGACC-3' |
| *Hyal1* | 5’-CATGCCTGAACCTGACTTCT-3’ | 5’-GTAGCAGTCAGGGAAGCCATA-3’ |
| *Hyal2* | 5’-CACCTGCCCATGCTGAAGGA-3’ | 5’-TCAGGAAAGAGGTAGAAGCC-3’ |
